# Identification of different classes of genome instability suppressor genes through analysis of DNA damage response markers

**DOI:** 10.1101/2022.02.28.482315

**Authors:** Bin-Zhong Li, Richard D. Kolodner, Christopher D. Putnam

## Abstract

Genetic studies in *Saccharomyces cerevisiae* have identified 266 genes and predicted an additional 38 genes that suppress the accumulation of spontaneous gross chromosomal rearrangements (GCRs). Here we identified mutations that induce two DNA damage response (DDR) markers, Hug1 expression and Ddc2 foci, and combined these data with other published screens identifying mutations inducing other DDR markers, including Rad52 foci and Rnr3 expression. Together, these data identify four categories of mutations: most mutations were DDR- GCR-, 356 were DDR+ GCR-, 72 were DDR- GCR+, and 157 were DDR+ GCR+. These results indicate that induction of DDR markers alone, while allowing DDR analysis, is not a reliable indicator of increased genome instability. They also guide further analysis of mechanistically distinct groups of GCR-inducing mutations, such as mutations that increase levels of GCR-inducing DNA damage and mutations that result in mis-repair of normal levels of DNA damage resulting in GCRs.

## INTRODUCTION

Cells have sensitive mechanisms to identify and signal the presence of DNA damage. Mutations affecting these pathways have been linked to cancer predisposition syndromes in humans and increased genome instability in both model organisms and human cells and tumors (Carbone et al., 2020; Friedberg et al., 2013; Sharma et al., 2020; Vogelstein and Kinzler, 2004). Markers for the presence of DNA damage have been widely exploited as *in vivo* sensors for increased levels of DNA damage (Downs et al., 2000; Fernandez-Capetillo et al., 2002; Lou et al., 2003; Maser et al., 1997; Melo et al., 2001; Nelms et al., 1998; Paull et al., 2000; Stewart et al., 2003; Sun et al., 1996; Wang et al., 2002; Zhong et al., 1999). In spite of their utility in detecting signaling of the presence of DNA damage, it has not been clear if increased levels of DNA damage markers are reliable markers of increased genome instability, such as increased accumulation of gross chromosomal rearrangements (GCRs).

In *S. cerevisiae*, many markers indicating the presence of DNA damage have been investigated. A number of these markers detect increased phosphorylation due to DNA damage signaling by the protein kinase cascade comprised of Tel1^ATM^ or Mec1^ATR^- Ddc2^ATRIP^, Rad53^CHEK2^, and Dun1 or detect the consequences of the activation of this phosphorylation cascade (Ciccia and Elledge, 2010; Lanz et al., 2019; Putnam et al., 2009b). Markers of the activation of this pathway also include the accumulation of phosphorylated histone H2A (the *S. cerevisiae* equivalent of γ of Rad55, hyperphosphorylation of Rad53^CHEK2^, and the formation of Ddc2^ATRIP^ and Ddc1^RAD9A/RAD9B^ foci (Bashkirov et al., 2006; Downs et al., 2000; Melo et al., 2001; Sun et al., 1996). Moreover, activation of this phosphorylation cascade also leads to induction of a number of proteins encoded by ribonucleotide reductase (RNR)-related genes, including genes encoding some RNR subunits (*RNR2*, *RNR3*, and *RNR4*) and *HUG1*, which encodes a small protein inhibitor of RNR. Hug1, along with the related inhibitor proteins Dif1 and Sml1, controls RNR activity through nuclear sequestration of the Rnr2- Rnr4 heterodimer and inhibition of the Rnr1 homodimer (Desany et al., 1998; Huang et al., 1998; Lee and Elledge, 2006; Lee et al., 2008; Meurisse et al., 2014; Wu and Huang, 2008; Zhang et al., 2006; Zhao et al., 1998). This DNA damage checkpoint-induced transcription is mediated by hyperphosphorylation of Rfx1/Crt1, preventing its DNA binding and its recruitment of the general Tus1-Cyc8 co-repressor (Huang et al., 1998). Increased RNR activity is the essential function of the Mec1^ATR^-Ddc2^ATRIP^ and Rad53^CHEK2^ checkpoint proteins; deletion of the genes encoding any of the RNR inhibitors *DIF1*, *HUG1*, or *SML1* suppresses the lethality of the deletion of these checkpoint genes (Basrai et al., 1999; Desany et al., 1998; Lee et al., 2008; Wu and Huang, 2008; Zhao et al., 1998). A number of DNA repair proteins, such as Rad54 and RPA, are also targets of phosphorylation by the DNA damage signaling protein kinase cascade and this has also been used as a marker of DNA damage signaling; phosphorylation of DNA repair proteins can also alter their activity (Bashkirov et al., 2000; Chen et al., 2010; Herzberg et al., 2006; Lanz et al., 2021; Smolka et al., 2007).

Many proteins form discrete nuclear foci in response to DNA damage, most notably proteins that function in double strand break (DSB) repair (Lisby et al., 2004; Lisby et al., 2003; Lisby and Rothstein, 2009; Maser et al., 1997; Nelms et al., 1998) but also other types of DNA repair proteins and checkpoint proteins that signal the presence of DNA damage (Hombauer et al., 2011; Makhnevych et al., 2009; Melo et al., 2001; Tkach et al., 2012; Yimit et al., 2015). For example, nuclear foci formed in response to DSBs have been identified for the Mre11-Rad52-Xrs2 (MRX) complex, Rad52, Rad51, Rad54, Rad55, Rad59, Rdh54 and RPA and these co-localize to the sites of DSBs as well as with checkpoint proteins such as Ddc2 that assemble at DSBs as part of the checkpoint activation pathways (Lisby and Rothstein, 2009; Maser et al., 1997; Melo et al., 2001; Nelms et al., 1998). In addition, genetic studies using foci formation as a readout in *S. cerevisiae* have elucidated an ordered assembly pathway for DSB repair proteins in which RPA first binds to single stranded regions of DNA at the DSB followed by recruitment of Rad52 which then initiates the recruitment of the other DSB repair proteins Rad59 and Rad51, with Rad51 recruiting Rdh54, Rad55/57 and then Rad54 (Lisby et al., 2004).

We have previously performed multiple genetic screens of *S. cerevisiae* mutation collections to identify mutations causing increased GCR rates. These screens identified 266 non-essential and essential genes in which mutations cause increased rates of accumulating spontaneous GCRs and predicted an additional 38 such genes (Putnam et al., 2012; Putnam et al., 2016; Srivatsan et al., 2019). In addition, screens for mutations giving rise to a variety of other types of genome instability including chromosomal instability, direct repeat instability, and loss-of-heterozygosity have been performed in other laboratories (Andersen et al., 2008; Novarina et al., 2020; Stirling et al., 2011; Yuen et al., 2007). Together these datasets provide an opportunity for us to evaluate how well markers of increased DNA damage signaling correlate with increased genome instability. The data from DNA damage signaling marker screens used in our analysis include those for Rnr3 induction and Rad52 foci formation (Alvaro et al., 2007; Hendry et al., 2015; Styles et al., 2016) as well as a whole-genome screen for Hug1 induction and a focused screen for Ddc2 foci formation performed for this study. The analyses reveal that many mutations cause differential responses, depending on the DNA damage marker analyzed. Many of the mutations cause either (1) high damage signaling and high genome instability or (2) low damage signaling and low genome instability, which is consistent with a correlation of DNA damage levels and genome instability. In addition, we identified many other mutations that only cause increased damage signaling or only increased genome instability. Together, our results argue for caution in inferring the presence or lack of increased genome instability solely based on the presence or lack of markers for increased DNA damage, particularly if only a single DNA damage marker is tested.

## RESULTS

### Selection of a marker for monitoring the DNA damage response

We first analyzed four genes whose expression was substantially increased (*RNR3*, *DDR2*, and *HUG1*) or decreased (*DSE2*) by induction of DNA damage (Greenall et al., 2008). These genes were analyzed by generating four strains in which one of the genes was tagged with *EGFP*. The resulting strains were grown to mid-log phase and treated with and without 100 mM hydroxyurea for 2 hours to induce DNA damage, and the EGFP signal was measured by fluorescence-activated cell sorting (FACS). Because FACS analysis determines the EGFP signal for each cell, this method is even suitable for analysis of strains that have a slow growth phenotype. We found that the protein levels of Rnr3-EGFP and Hug1-EGFP were substantially up-regulated by HU treatment, which is consistent with the results of previous studies (Basrai et al., 1999; Huang et al., 1998), but Dse2-EGFP or Ddr2-EGFP were not substantially changed (**Suppl. Fig. 1A**). Because Hug1-EGFP showed greater induction, we selected the *HUG1-EGFP* fusion as a marker for detecting induction of DNA damage *in vivo*. Previous studies have similarly used a reporter consisting of GFP or firefly luciferase driven by the *HUG1* promoter to show that Hug1 expression is sensitive to a wide variety of DNA damaging agents but not other types of cellular stress (Ainsworth et al., 2012; Benton et al., 2007).

### Identification of genes suppressing Hug1-EGFP induction

To identify mutations causing elevated levels of Hug1 induction in the absence of exogenous DNA damaging agents, we first introduced the *HUG1-EGFP* fusion gene linked to a *HIS3* marker into a MATα bait strain. We then crossed this bait strain to the BY4741 **MATa** collection of *S. cerevisiae* strains with deletions of all non-essential genes using a modified synthetic genetic array (SGA) protocol (Putnam et al., 2016; Tong and Boone, 2006). A control strain that could be selected for by SGA was generated by replacing the *leu2*Δ*0* allele in BY4741 with a *leu2*Δ*::kanMX4* allele. Our SGA protocol selected for **MATa** haploids containing *HUG1-EGFP* and the gene deletion of interest from the deletion collection strain. Unperturbed log-phase cultures were analyzed for DNA content by high-throughput FACS; strains that did not have a haploid DNA content or contained a mixture of cells with haploid and diploid DNA content were eliminated from consideration. The unperturbed log-phase cultures were also analyzed by high-throughput FACS to determine the levels of EGFP fluorescence (**Fig. 1A**). To determine the level of Hug1-EGFP induction, we determined the average EGFP signal for each strain and normalized these averages to that of control strains analyzed at the same time to determine the fold induction of Hug1-EGFP signal for each strain (**Suppl. Table 1**). Because subsets of the deletion collection were independently mated and analyzed up to three times, 4,792 deletion mutants derived from 5,781 independently generated strains with haploid DNA content were analyzed; repeated measurements from independent biological isolates were highly correlated (**Suppl. Fig. 1B**).

**Figure 1.**
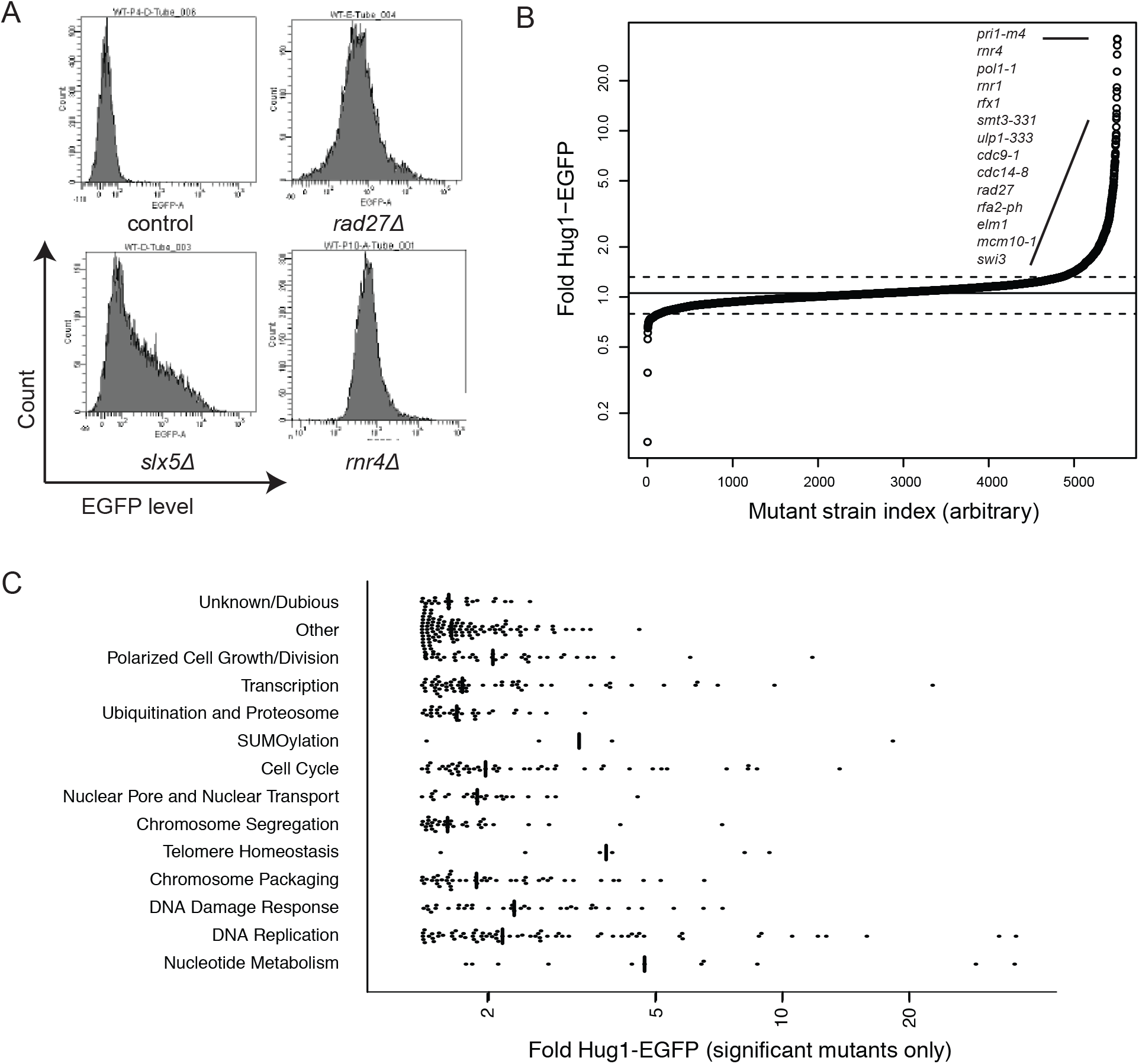
Distribution of Hug1-EGFP levels in the mutants analyzed. **A. Histograms** of Hug1-EGFP levels in individual cells from log phase cultures of the control (*leu2*Δ) strain and three representative mutant strains having increased levels of Hug1-EGFP expression. **B.** Distribution of the fold Hug1-EGFP levels for all of the deletion and temperature sensitive mutants analyzed. The horizontal line indicates the center of the fitted normal distribution (**Suppl. Fig. 1C**). **C.** Beehive plot of the fold increase in Hug1-EGPF levels for the significant mutants broken down by functional category.

The Hug1-EGFP expression data for 4,792 deletion mutations was combined with our previously published Hug1-EGFP expression data for 394 strains containing temperature sensitive mutations (**Fig. 1B**; (Srivatsan et al., 2019)). The resulting fold increased Hug1-EGFP distribution was normally distributed with a mean of 1.06 and a standard deviation of 0.13 (**Suppl. Fig. 1C**). This result is consistent with the hypothesis that most mutations do not substantially alter Hug1 expression. The experimental distribution had more strains with increased Hug1 expression than would be predicted from a normal distribution (**Suppl. Fig. 1C,D**); these strains have mutations that induce Hug1 expression during unperturbed growth.

Using a false-discovery rate of 0.05, 479 mutations affecting 416 genes were identified as causing increased levels of Hug1-EGFP expression, with a fold increased Hug1-EGFP expression range of 1.39 to 35.8. Many of the genes identified act in DNA metabolism-related pathways: nucleotide metabolism, DNA replication, DNA repair, post-replicative chromosome packaging (chromatin assembly, sister chromatid cohesion, chromosome condensation), chromosome segregation, polarized cell growth, and cell cycle progression (**Fig. 1C**; **Suppl. Table 1**). In addition, mutations in genes involved in maintaining telomere homeostasis, the nuclear pore and nuclear import/export processes, protein sumolyation, protein degradation, the vacuolar ATPase (V-ATPase), a Golgi mannosyltransferase, TOR signaling, mRNA nonsense mediated decay, and the mRNA cleavage and polyadenylation factor 1A also increased Hug1 expression (**Suppl. Table 1**).

### A subset of mutations that induce Hug1 expression also induce Rnr3 expression

Because of the similarities of *RNR3* and *HUG1* regulation in response to DNA damage (Basrai et al., 1999; Huang et al., 1998), we anticipated that Hug1 induction would show high correlation with previous measurements of Rnr3-GFP induction as assayed by colony fluorescence (Hendry et al., 2015). Of the 4,467 mutations tested in both assays, 73 mutations in 67 genes caused increased DNA damage signaling in both assays (**Fig. 2; Suppl. Table 2**). In contrast, 317 mutations in 274 genes caused significant induction in only the Hug1 assay, and 61 mutations in 61 genes caused significant induction in only the Rnr3 assay (**Fig. 2**). The 73 mutations identified using both assays tended to affect genes involved in DNA replication (*CDC8, DBF4, DNA2, MCM10, MRC1, ORC1, ORC2, POL1, POL12, POL2, POL3, POL31, POL32, PRI1, PRI2,* and *RFA2*), DNA damage response and repair (*DIA2, ESC2, MMS1, MMS22, MRE11, NSE1, RAD5, RAD51, RAD52, RAD54, RAD55, RMI1, RTT101, RTT107, SGS1, SLX5, SLX8,* and *TOP3*), and sumoylation (*SMT3* and *ULP1*). Consistent with previous results (Basrai et al., 1999), mutations affecting the mechanism that normally suppresses the transcriptional response to DNA damage, including *rfx1Δ/crt1Δ*, *isw2Δ*, *itc1Δ*, and *hda3Δ*, caused increased expression of both Hug1 and Rnr3, with loss of *RFX1/CRT1* causing the greatest induction. Other mutations known to affect genes involved in the transcriptional repression of *HUG1* and *RNR3* were either not tested (*TUP1* and *CYC8/SSN6*) or were only significant in a single assay (*HDA1* and *SIN4*).

**Figure 2.**
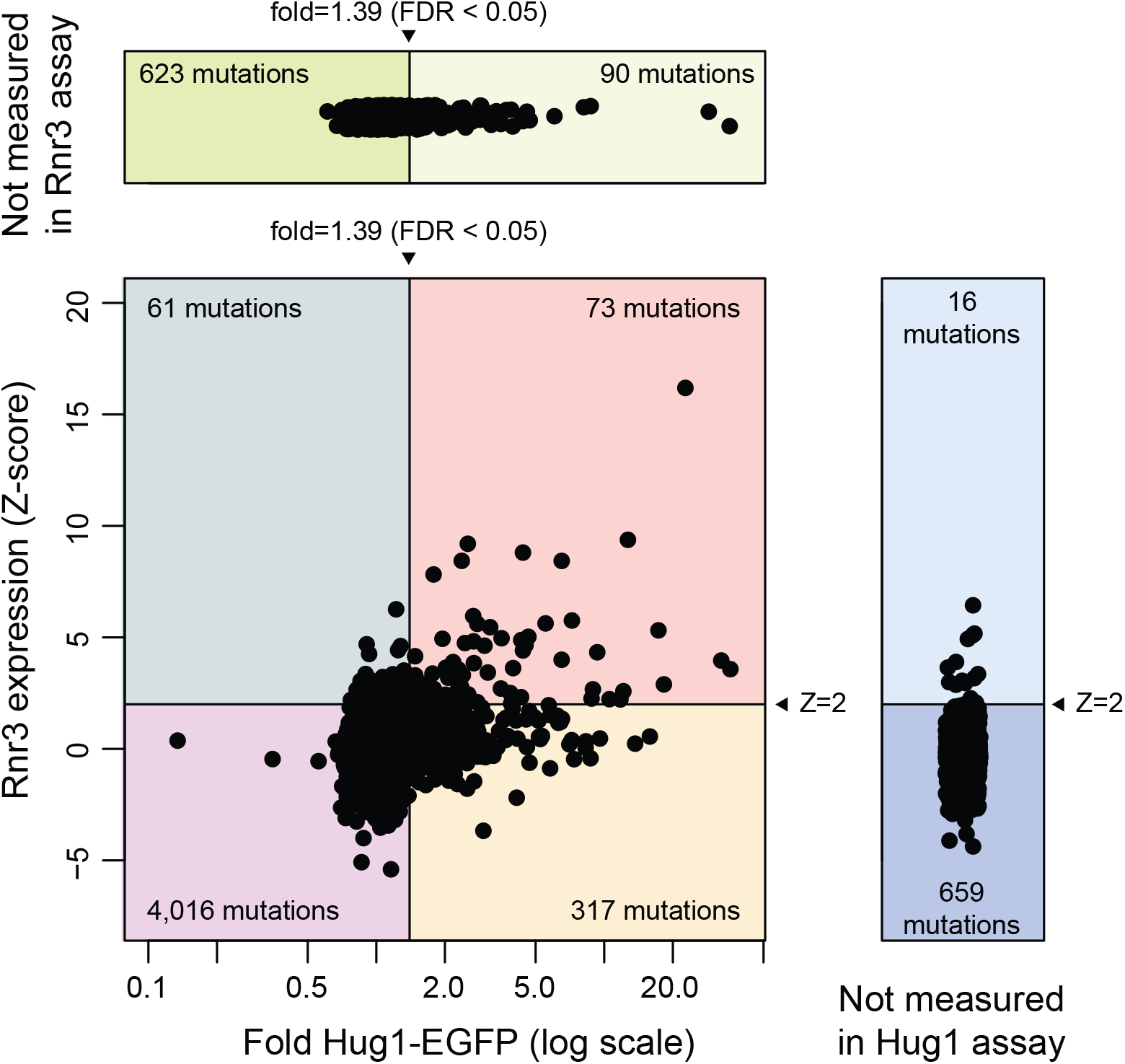
Comparison of Hug1 and Rnr3 induction in the mutants analyzed. Scatter plot of the Rnr3 expression Z-score (Hendry et al., 2015) vs. the fold increase in Hug1- EGFP expression for the mutants analyzed in common. Horizontal and vertical lines are the threshold of significance for the Rnr3 and Hug1 expression data. Beehive plots above and to the right illustrate the distribution of the fold Hug1-EGFP expression and the Rnr3 expression for mutants tested in only one of the two assays.

### Induction of Ddc2 foci imperfectly correlates with Hug1/Rnr3 induction

We next explored the correlation of Hug1 and Rnr3 induction with the accumulation of Ddc2 foci, which is a marker of sites of DNA damage and checkpoint activation (Melo et al., 2001; Waterman et al., 2019). We constructed a **MAT**α bait strain expressing a Ddc2-EGFP fusion protein and the nuclear pore fusion protein Nup49-mCherry, which allows visualization of the nuclear envelope to ensure that only nuclear foci were scored. We crossed this bait strain using SGA to 469 BY4741 **MATa** strains representing deletion mutations that induced Hug1 expression and/or increased GCR rates, some of which also induced Rnr3 expression. The resulting haploid strains were grown to log phase in complete synthetic media and were analyzed by fluorescence microscopy to determine the fraction of cells containing nuclear Ddc2 foci (**Fig. 3A; Suppl. Table 3**). We assessed significance thresholds for the fraction of cells with Ddc2 foci using the 5^th^ percentile (fraction of cells with Ddc2 foci=0.015) and 95^th^ percentile (fraction=0.048) of the observations of the *leu2Δ::kanMX4* control strain (average=0.031, standard deviation=0.013). Only 4 of the 444 mutant strains recovered had levels of Ddc2 foci below the lower threshold: *npl3*Δ, *ydjl*Δ, *dus1*Δ, and *mec1*Δ *sml1*Δ. In contrast, 153 mutant strains with defects in 152 genes had levels of Ddc2 foci that were above the upper threshold (**Fig. 3B; Suppl. Table 3**).

**Figure 3.**
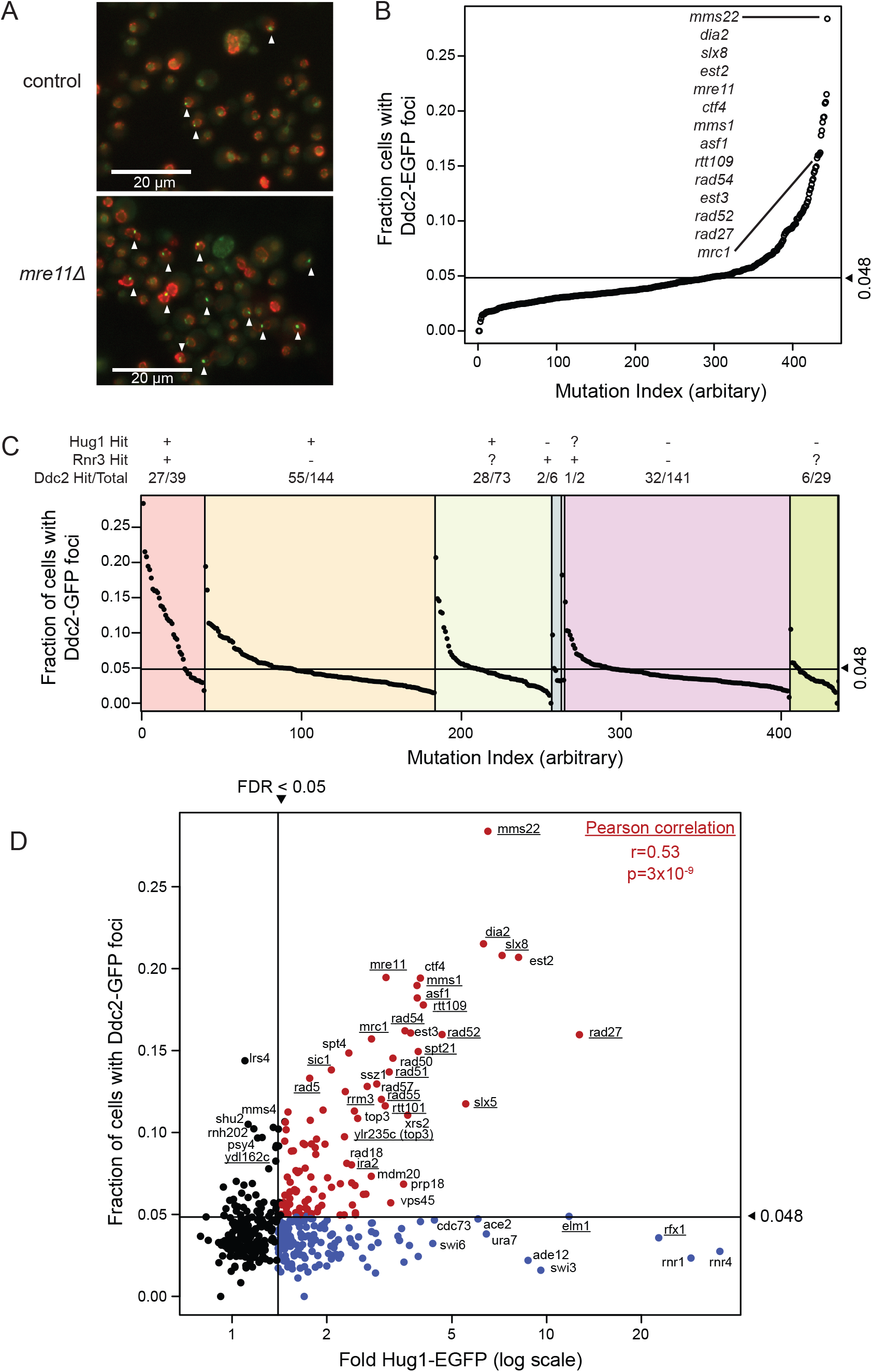
Distribution of the fraction of cells with Ddc2-EGFP foci. **A.** Representative images of Ddc2-EGFP foci for a control (*leu2*Δ) and an *mre11*Δ strain. **B.** Distribution of the fraction of cells with Ddc2-EGFP foci. The horizontal line at a fraction of 0.048 is the 95% percentile for the *leu2*Δ control strain measurements. **C.** Distribution of the fraction of cells with Ddc2-EGFP foci grouped by the effect of the mutations in the Hug1 and Rnr3 expression assays; “+” indicates a significant effect in that assay, “-” indicates not significant, and “?” indicates not measured. The fraction above each category indicates the number of mutations causing a significant increase in Ddc2 foci over the total number of mutations in that category. **D.** Scatter plot of the fraction of cells with Ddc2-EGFP foci vs. the fold Hug1-EGFP expression for each mutant analyzed. Red points indicate mutants with increased Hug1-EGFP expression that correlate with increased levels of Ddc2-EGFP foci, and blue points indicate mutants with increased Hug1-EGFP expression that does not correlate with increased levels of Ddc2-EGFP foci. The horizontal and vertical lines indicate the significance threshold for each assay. The Pearson correlation coefficient (*r*) and *p*-value for the null model of the correlation being zero were calculated using only the points in the upper righthand quadrant. Underlined mutants had a significant increase in Rnr3 expression.

Mutations causing increased expression of both Hug1 and Rnr3 tended to cause increased levels of Ddc2 foci levels (27 of 39 mutations; **Fig. 3C**), consistent with a role of Mec1-Ddc2 in the signaling cascade leading to Hug1 and Rnr3 induction (Basrai et al., 1999; Huang et al., 1998). The mutations that caused increased expression of both Hug1 and Rnr3 and increased levels of Ddc2 foci primarily affected genes encoding proteins that function in DSB repair, DNA replication and replication fork processing (*DIA2*, *ELM1*, *ESC2*, *IRA2*, *MMS1*, *MMS22*, *MRC1*, *MRE11*, *POL32*, *RAD5*, *RAD27*, *RAD51*, RAD52, *RAD54*, *RAD55*, *RMI1*, *RRM3*, *RTT101*, *RTT107*, *RTT109*, *SGS1*, *SIC1*, *SLX5*, *SLX8*, *SPT21*, *TOP3/YLR235C*, and *WSS1*; **Fig. 3C**). Mutations that induced both Hug1 and Rnr3 expression but did not increase Ddc2 foci affected genes involved in transcriptional repression of *HUG1* and *RNR3* (*HDA3*, *ISW2*, *ITC1*, and *RFX1*) as well as genes not clearly linked to DNA replication or the DNA damage response (*ARV1*, *ELM1*, *ETR1*, *GAS1*, *LEO1*, *LSM6, RTF1*, and *SAC3*). Conversely, mutations causing increased Ddc2 foci also were not only observed in the set of mutations that induced both Hug1 and Rnr3, but were also observed in the sets of mutations that only induced Hug1, only induced Rnr3, and did not induce either Hug1 or Rnr3 (**Fig. 3C**). Some of the mutations that did not induce Hug1 or Rnr3 but caused increased Ddc2 foci affected genes involved in checkpoint activation (*RAD24*, *RAD17*, *DDC1*, and *RAD9*), genes which would be expected to be required for Hug1 and Rnr3 induction. Other mutations only identified in the Ddc2 foci screen included those in genes encoding RNase H2 subunits (*RNH202* and *RNH203*) and in a number of other genes with roles in DNA replication, repair and damage signaling (*APN1*, *CSM3*, *DPB3*, *MMS2*, *MMS4*, *MPH1*, *PSY4*, *SHU2*, *SIZ1*, and *UBC13*); these mutations were included in the screen due to the fact that they increased GCR rates either as single mutants or in combination with other mutations.

Direct comparison of Hug1 induction with Ddc2 foci levels revealed that Hug1-inducing mutations belonged to two groups (**Fig. 3D**). The first group contained 110 mutations in which the increased levels of Ddc2 foci were correlated to the increased levels of Hug1 expression (**Fig. 3D**, upper right quadrant). This is consistent with these mutations causing the accumulation of different levels of DNA damage, leading to Mec1-Ddc2 signaling and Hug1 induction. This first group included many mutations affecting DNA damage response and avoidance genes (*SLX5*, *SLX8*, *MRE11*, *RAD50*, *XRS2*, *RAD51*, *RAD52*, *RAD54*, *RAD55*, *RAD57*, *RAD5*, *RAD18*, *RAD27*, *MMS1*, *RTT109*, *RTT101*, *SGS1*, *TOP3*, *RMI1*, *SIC1*, *SPT4*, *ASF1*, *CTF4*, *MMS22*, and *DIA2*). The second group contained 146 mutations that caused Hug1 induction but not increased Ddc2 foci (**Fig. 3D**, lower right quadrant). Given the role of Mec1-Ddc2 signaling in DNA damage-induced Hug1 induction (Basrai et al., 1999), these mutations seem likely to affect Hug1 induction independently of Mec1-Ddc2 DNA damage signaling, such as through loss of the *HUG1* gene regulation (such as deletions of *RFX1/CRT1*, *ITC1*, and *ISW1*), through transcriptional responses due to changes in nucleotide pools (such as deletions in *RNR1*, *RNR4*, *ADE12*, and *URA7*), or through changes in other processes that may indirectly affect Hug1 expression (such as deletions in *ELM1* and *SWI3*).

### Rad52 foci induction shows poor correlation with the Hug1/Rnr3 transcriptional response

We also compared the effect of mutations on Hug1 and Rnr3 induction with their effect on the accumulation of Rad52-YFP foci in uninduced diploid cells, previously called the Increased Recombination Center (IRC) phenotype (Alvaro et al., 2007). We note that a second genome-wide Rad52 foci screen has been performed in haploid strains (Styles et al., 2016); however, as only the list of mutations causing increased Rad52 foci in haploid strains were reported, comparisons with the haploid screen will be described separately below. Rad52 foci are homologous recombination (HR) repair intermediates that predominantly form during S-phase where they are thought to play a role in replication fork progression rather than acting as a component of DNA damage signaling (Lisby et al., 2001); Rad52 foci are also thought to act in DSB repair. As with Ddc2 foci, increased levels of Rad52 foci in the diploid screen were caused by mutations that caused increased Hug1 and Rnr3 expression as well as by mutations that did not increase either Hug1 or Rnr3 expression (**Fig. 4A**). Hence, increased induction of Hug1 and Rnr3 expression were imperfect predictors of the likelihood that a mutation causes increased levels of Rad52 foci. Mutations identified in both the Hug1 and Rnr3 induction assays included the highest percentage of mutations causing increased Rad52 foci (41%), whereas the percentage of mutations causing increased Rad52 foci was much lower for mutations identified in only one of the Hug1 or Rnr3 expression assays (8%) or not identified in either assay (4%). However, given the number of mutations in the latter categories, the number of mutations that caused increased Rad52 foci and were identified both of the Hug1 and Rnr3 assays (16 mutations) is smaller than those causing increased Rad52 foci that were identified in only one of the Hug1 and Rnr3 expression assays (22 mutations) or were not identified in either of the Hug1 or Rnr3 expression assays (150 mutations). The 16 mutations that resulted in increased induction of Hug1 and Rnr3 expression and increased Rad52 foci all also resulted in increased Ddc2 foci and primarily affected genes encoding proteins that function in DSB repair, DNA replication and replication fork (*ESC2*, *MMS1*, *MRE11*, *POL32*, *RAD27*, *RAD51*, *RAD54*, *RAD55*, *RMI1*, *RRM3*, *RTT101*, *RTT107*, *RTT109*, *SGS1*, *SLX8*, and *WSS1*). Moreover, the level of induction seen in the Hug1 and Rnr3 assays was not predictive of the levels of induction of Rad52 foci; some of the mutations causing the highest levels of Rad52 foci were observed in strains with mutations that did not increase either Hug1 or Rnr3 expression (**Fig. 4A**).

**Figure 4.**
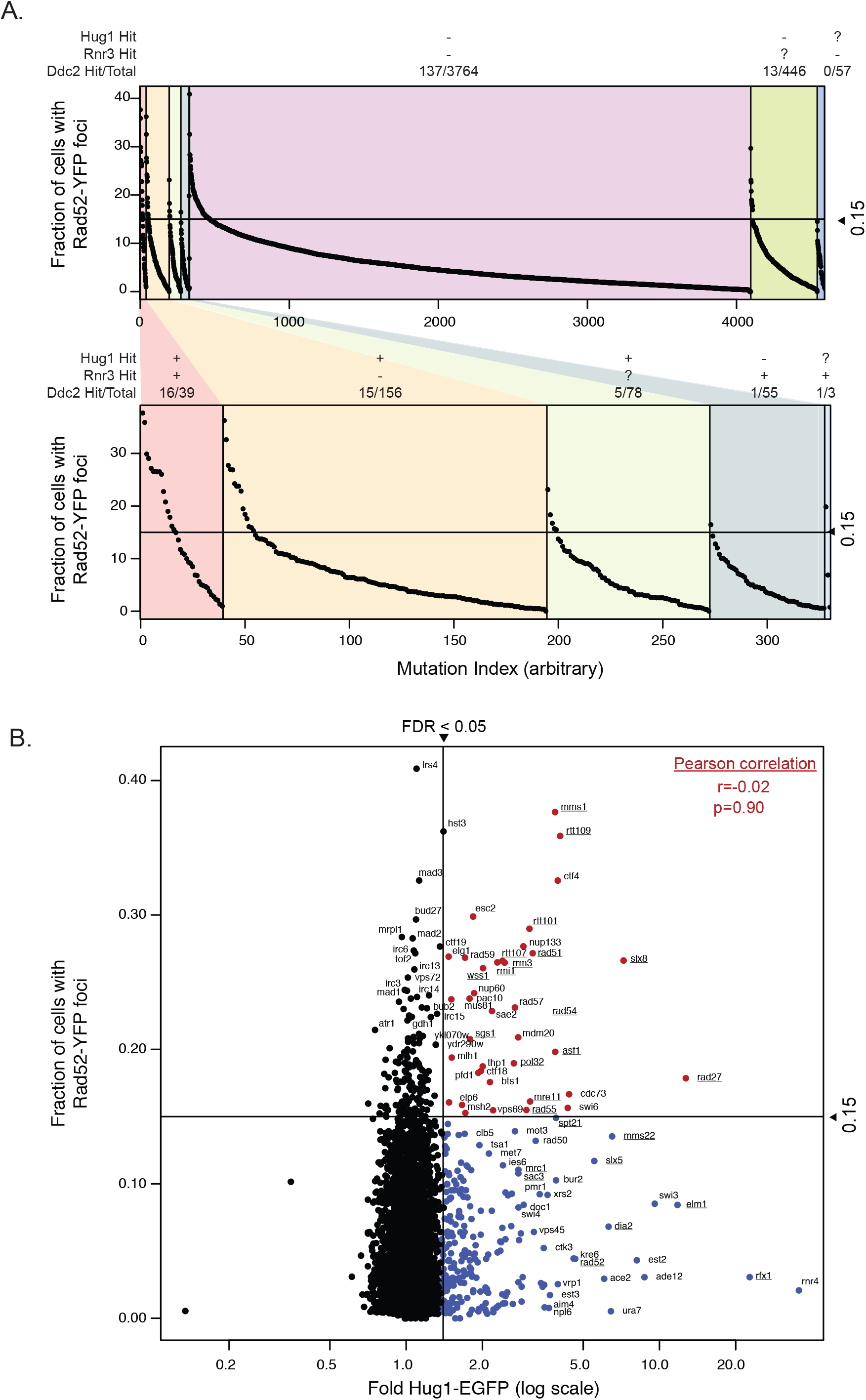
Distribution of the fraction of cells with Rad52-YFP foci. **A.** Distribution of the percent of cells with Rad52-YFP foci (Alvaro et al., 2007) grouped by the effect of the mutations on Hug1 and Rnr3 expression; “+” indicates a significant effect in that assay, “-” indicates not significant, and “?” indicates not measured. The fraction above each category indicates the number of mutations causing a significant increase in Rad52 foci over the total number of mutations in that category. **B.** Scatter plot of the fraction of cells with Rad52-YFP foci vs. the fold Hug1-EGFP expression for each mutant analyzed. Horizontal and vertical lines indicate the significance threshold for each assay. The Pearson correlation coefficient (*r*) and *p*-value for the null model of the correlation being zero were calculated using only the points in the upper righthand quadrant. Underlined mutants had a significant increase in Rnr3 expression.

We found that there was limited if any correlation between the amount of Hug1 induction and Rad52 foci induction caused by mutations that induced both Hug1 expression and Rad52 foci (**Fig. 4B**). This difference from Ddc2 foci induction is likely due to the fact that Rad52, unlike Ddc2, is not directly in the Hug1/Rnr3 signal transduction pathway and that the kinds of damage that induce Rad52 foci may only partially overlap the kinds of damage that induce the Hug1/Rnr3 signal transduction pathway. In addition, increased Rad52 foci could be caused by defects resulting in (1) increased DNA damage, (2) increased processing of normal levels of endogenous DNA damage by HR, and (3) delays in the turnover of HR intermediates. The possibility that both direct and indirect factors contribute to accumulation of Rad52 foci is echoed by the diverse biological roles of the genes identified as suppressing the IRC phenotype (Alvaro et al., 2007), including transcription initiation (*IRC1*), mitochondrial genome maintenance (*IRC3*), clathrin mediated-vesicle trafficking (*IRC6*), the degradation of cysteine to ammonia and pyruvate (*IRC7*), proteasome assembly (*IRC25*), and synthesis-dependent DNA strand annealing (*IRC20*).

### Alterations in cell cycle distribution do not account for the induction of DDR markers

Induction of DDR markers can occur at different stages of cell cycle progression, and hence cell cycle distribution differences could contribute to DDR marker induction (Escribano-Diaz et al., 2013; Jazayeri et al., 2006; Meurisse et al., 2014). We therefore performed an extensive analysis of published data (**Suppl. Appendix 1**) to examine the cell cycle distribution of unperturbed log-phase cells generated by FACS analysis of the *S. cerevisiae* haploid (Koren et al., 2010; Soifer and Barkai, 2014) and diploid (Hoose et al., 2012) deletion collections (**Suppl. Table 4**). Each measured cell cycle distribution (**Fig. 5A**) was fitted so that the percentage of cells in the G1-, S-, and G2-phases of the cell cycle summed to 100% (**Fig. 5B**), and was plotted as a single point on a ternary plot diagram (**Fig. 5C**). The percentage of G1-, S-, and G2-phase cells was 21.5%, 25.1%, and 53.4% for the haploid control (n=292), and 22.8%, 24.3%, and 52.3% for the diploid control (n=219, **Suppl. Fig. 2**). Mutations causing substantially altered cell cycle distributions were also identified (**Fig. 5D, Suppl. Appendix 1, Suppl. Table 4**).

**Figure 5.**
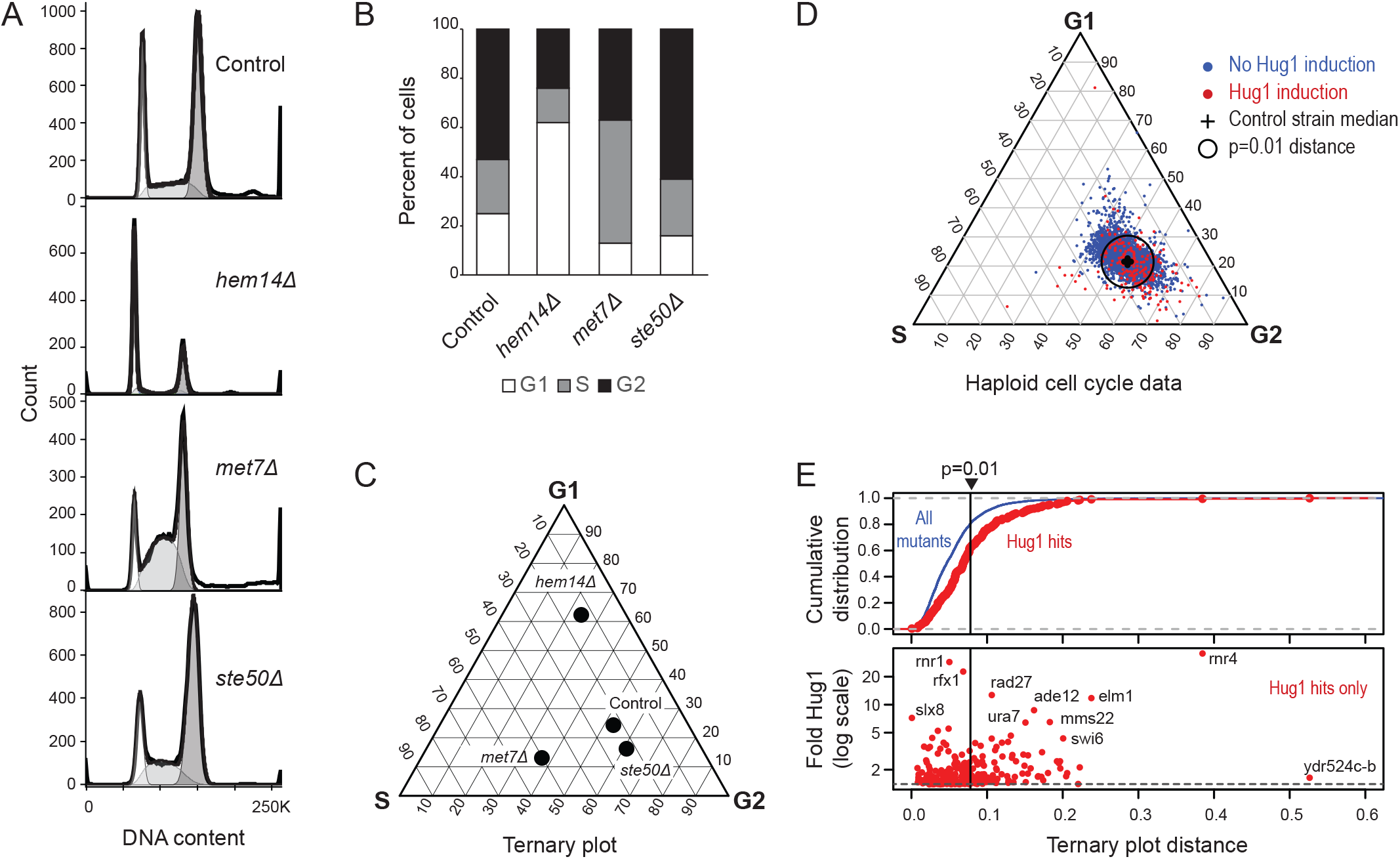
Cell cycle distributions for mutant strains with and without increased levels of DNA damage markers. **A.** Example FACS analyses of log-phase cultures of a haploid *S. cerevisiae* control strain and haploid strains with enriched G1 (*hem14*Δ), S (*met7*Δ), and G2 (*ste50*Δ) populations. FACS data (Koren et al., 2010) were plotted using FlowJo (FlowJo, 2019) with cell count on the y-axis and the measured SYBR-green area for each cell on the x-axis. Peaks for these haploid strains are at 1n and 2n DNA content. **B.** Stacked bar plots for the cell cycle distributions for the observations in panel A derived using FlowJo with the Dean-Jett-Fox model (Fox, 1980) and fitted to account for cell-cell aggregation. **C.** Ternary plot of the cell-cycle distributions of the observations in panel A. Strains consisting of 100% G1 cells would be at the top vertex, those with 100% S cells would be at left vertex, and those with 100% G2 cells would be at the right vertex. **D.** Ternary plot diagram for the cell cycle distribution of haploid strains colored by the effect of the mutation in these strains on Hug1 induction (blue=no induction, red=induction). The black cross indicates the position of the wild-type control strain. Strains indicated outside the black circle have a significantly altered cell cycle distribution (p<0.01) compared to the wild-type control strain. **E.** Analysis of Hug1 induction of mutations based on the ternary plot distance between the mutant distribution and the geometric median of the control strain distribution. (Top) The cumulative distribution of all mutations (blue) is compared the cumulative distribution of distances for those mutations with significant induction of Hug1. (Bottom) Plot of the fold Hug1 induction against the ternary plot distance. The vertical and horizontal lines indicate significance cutoffs for the two assays.

The cell cycle distributions were compared for mutations inducing DDR markers (DDR+ mutations) and mutations not inducing DDR markers (DDR-mutations). Most mutations inducing Hug1 expression did not cause significantly altered haploid cell cycle distributions (in the p>0.01 ternary plot distances; **Fig. 5D, E**). However, when analyzed as a group, the Hug1-inducing mutations had greater alterations in their cell cycle distributions than mutations that did not induce Hug1 (p=3×10^-13^ Kolmogorov-Smirnov test; **Fig. 5E**), consistent with a fraction of mutations causing both Hug1 induction and an altered cell cycle distribution. The relative ability of DDR markers to predict cell cycle alterations was Hug1 induction > Ddc2 foci induction > Rnr3 induction > Rad52 foci induction, with the caveat that the Ddc2 foci induction data was generated from a selected set of mutations likely to bias its ranking in these results (**Suppl. Fig. 3**). Similarly, DDR+ mutations were more likely to have an altered cell cycle distribution in the haploid data than in the diploid data (**Suppl. Fig. 3-4**). Despite the results of this grouped analysis, only a subset of DDR+ mutations significantly altered cell cycle distributions, the magnitude of these alterations was not correlated with the DDR marker levels, and the magnitude of these alterations did not increase for mutations identified in increasing number of DDR marker screens (**Fig. 5E, Suppl. Fig. 5**). These results are consistent with (1) the ability of sufficiently high levels of DNA damage checkpoint activation to alter the timing of the cell cycle, (2) the likelihood that DDR markers are more sensitive probes for DNA damage than altered cell cycle distributions, and (3) the fact that DDR marker induction by most DDR+ mutations cannot be readily explained by altered cell cycle timing.

### Mutations causing increased GCR rates are not restricted to those causing increased DNA damage signaling

Extensive datasets of the effect of mutations in essential and non-essential *S. cerevisiae* genes on the accumulation of GCRs assessed in multiple GCR assays are now available (Chan and Kolodner, 2011; Chen and Kolodner, 1999; Nene et al., 2018; Putnam et al., 2009a; Putnam et al., 2016; Srivatsan et al., 2019). Because these GCR assays and the assays for the induction of DNA damage response markers were typically performed in the absence of exogeneous DNA damaging agents, these datasets can be directly compared. We initially observed that mutations causing increased GCR rates fell in groups of mutations that both did and did not cause Hug1 and Rnr3 induction (**Suppl. Fig. 6**). This analysis was then extended to include data generated with a broader set of DDR markers including Hug1 induction, Rnr3 induction (Hendry et al., 2015), increased Ddc2 foci, and increased Rad52 foci in diploids (Rad52-D; (Alvaro et al., 2007)) and in haploids (Rad52-H; (Styles et al., 2016)). Mutations for which GCR rates or patch scores were available were divided into two GCR-based groups (GCR+, mutations causing increased GCR rates in at least one GCR assay; GCR-, mutations that did not cause increased GCR rates) and three DDR-based groups (DDR+, mutations causing an increase in at least one DDR marker assay; DDR-, mutations that did not cause an increase in any DDR marker assay; and DDR?, mutations not tested in any DDR marker assay) (**Suppl. Fig. 7; Suppl. Table 5**). Many mutations fell into the DDR+ GCR+ (157 mutations) and the DDR-GCR- (810 mutations) categories, consistent with some correlation between increases in DDR markers and increased GCR rates. Of the 27 mutations that increased expression of both Hug1 and Rnr3 and increased levels of Ddc2 foci, 23 (85%) resulted in increased GCR rates and primarily affected genes encoding proteins that function in DSB repair, DNA replication and replication fork processing (*DIA2*, *ESC2*, *MMS1*, *MRC1*, *MRE11*, *POL32*, *RAD27*, *RAD5*, *RAD51*, *RAD52*, *RAD54*, *RAD55*, *RMI1*, *RRM3*, *RTT101*, *RTT107*, *SGS1*, *SIC1*, *SLX5*, *SLX8*, *SPT21*, *WSS1*); 16 of the 27 mutations also resulted in increased Rad52 foci and all 16 of the mutations that resulted in increased Rad52 foci also resulted in increased GCR rates (*ESC2*, *MMS1*, *MRE11*, *POL32*, *RAD27*, *RAD51*, *RAD54*, *RAD55*, *RMI1*, *RRM3*, *RTT101*, *RTT107*, *RTT109*, *SGS1*, *SLX8*, *WSS1*). Remarkably, we also observed many mutations in the DDR+ GCR- (356 mutations) and the DDR- GCR+ (72 mutations) categories, indicating that many genetic defects have a more complicated relationship between DNA damage/DNA damage responses and increased GCR rates; the genes implicated by the DDR- GCR+ fell into numerous classes including DNA replication initiation (*CDC7*, *DBF4*, *HCS1*, *ORC3*, *ORC5*, *RFA1*, *RFC5*), a subset of DNA repair genes including translesion synthesis (*DNL4*, *EXO1*, *HRQ1*, *OGG1*, *PIF1*, *RAD30*, *RDH54*, *REV1*, *REV3*, *REV7*, *RNH1*, *SHU1*, *SRS2*), the DNA damage checkpoint (*CHK1*, *DDC2*, *DUN1*, *MEC1*, *MEC3*, *MRC1*), the proteosome (*PRE10*, *RPN5*, *RPN10*, *RPT1*, *RPT2*, *SCL1*), the Rap1-Rif2 complex, sumoylation (*UBC9*), and cohesin, codensin, and the Smc5/6 complex (*IRR1*, *NSE4*, *NSE5*, *SMC2*, *SMC5*, *SMC6*, *YCG1*) (**Suppl. Table 5**). Importantly, the fold increase in GCR rates caused by many of the DDR- GCR+ mutations was as high or higher than the fold increase in GCR rates caused by the DDR+ GCR+ mutations that were associated with induction of increased levels of multiple DDR markers (**Suppl. Fig. 7**).

We also evaluated the mutations identified in whole-genome screens using other genome instability assays, including for the A-like faker (ALF) phenotype, the bi-allelic mating (BiM) phenotype, the chromosome transmission failure (CTF) phenotype, increased direct-repeat recombination (DR), and loss of heterozygosity (LOH) at the *MET15* and *SAM2* loci even though in most cases there is little information available about the types of genome rearrangements, if any, are caused by the mutations detected using these assays (Andersen et al., 2008; Novarina et al., 2020; Stirling et al., 2011; Yuen et al., 2007). The distribution of the mutations identified in these other assays was remarkably similar to that of the GCR+ mutations, and many of the mutations identified in these other genome instability assays belonged to the DDR- group of mutations (**Fig. 6**). Taken together, these results argue for caution in inferring the presence or lack of increased genome instability solely based on the presence or lack of markers for increased DNA damage.

**Figure 6.**
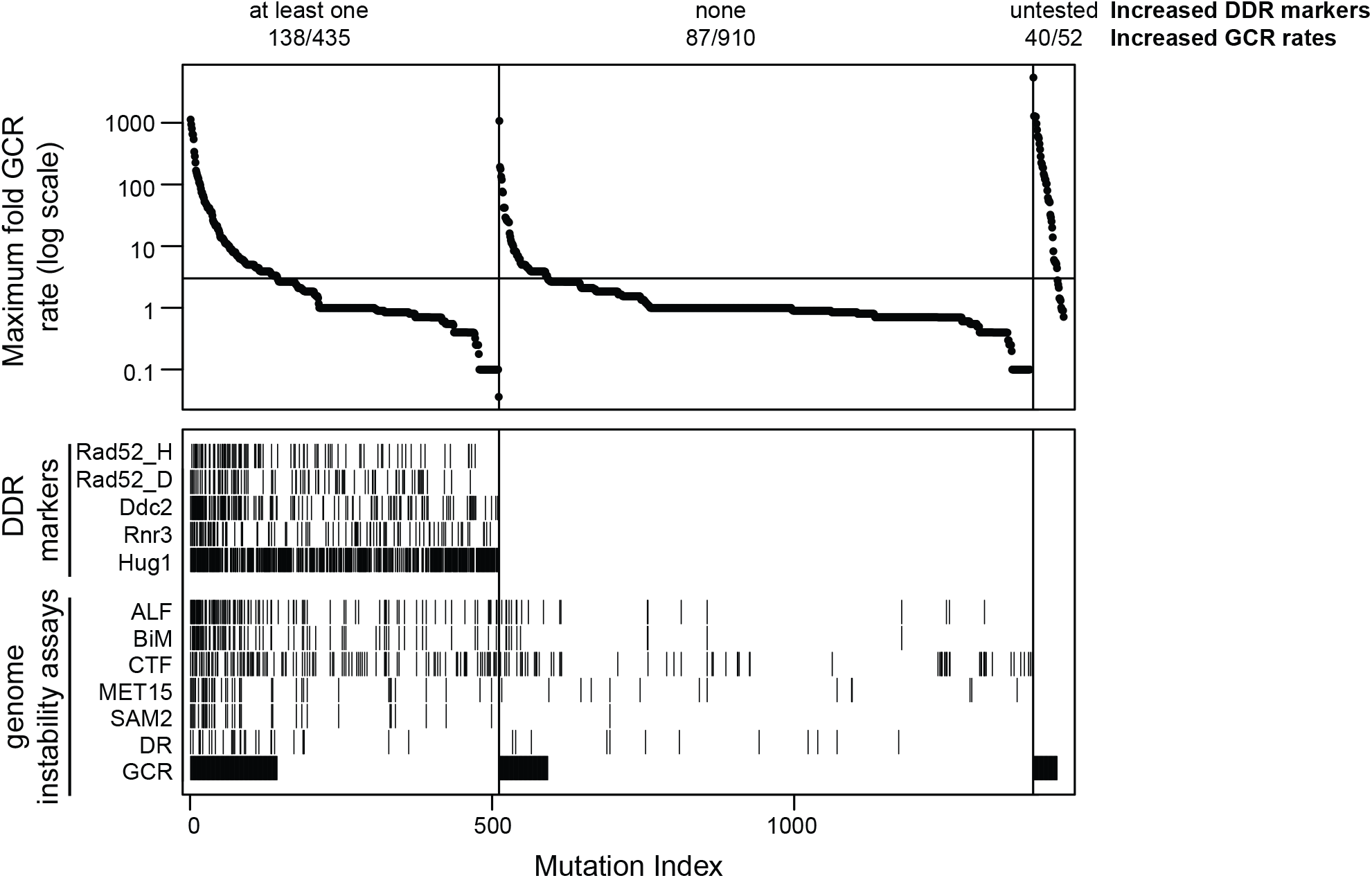
Distribution of the fold increase in GCR rate for the mutants analyzed. The distribution of maximum fold increase in GCR rates caused by each mutation in all of the GCR assays utilized are grouped by the effect of that mutation on Hug1 expression, Rnr3 expression, Ddc2 foci levels, and levels of Rad52 foci in haploid (Rad52_H) and diploid (Rad52_D) mutants. Groups of mutations are defined as those mutations causing an increase in at least one of the DDR marker assays, mutations without an increase in any DDR marker assay, and mutations not tested in any DDR marker assay. The fraction above each category indicates the number of mutations with a significant increase in GCR rates over the total number of mutations in that category. Vertical lines in the Rad52_H, Rad52_D, Ddc2, Rnr3, and Hug1 rows underneath the GCR rate distribution indicate that the mutation was found to cause a significant increase in that DDR marker assay. Vertical lines in the ALF, BiM, CTF, MET15 (MET15 LOH), SAM2 (SAM2 LOH), DR, and GCR rows indicate that the mutation was found be significant in the relevant genome instability assay. For this diagram, mutations with only patch scores in the duplication-mediated GCR assay (dGCR; (Putnam et al., 2016; Srivatsan et al., 2019)) but without a measured GCR rate were assigned an equivalent dGCR rate based on the relationship between the dGCR patch scores and dGCR rates (Putnam et al., 2016).

## DISCUSSION

Here, we evaluated the relationship between the ability of mutations to induce DDR markers and to induce genome instability. We performed screens to identify mutations causing increased levels of two DDR markers, expression of Hug1 and accumulation of Ddc2 foci. These data were aggregated with published data identifying mutations that increase levels of two other DDR markers, Rad52 foci and Rnr3 expression (Alvaro et al., 2007; Hendry et al., 2015; Styles et al., 2016), and that cause increased GCR rates and increased genome instability in other assays (Andersen et al., 2008; Chan and Kolodner, 2011; Chen and Kolodner, 1999; Nene et al., 2018; Novarina et al., 2020; Putnam et al., 2009a; Putnam et al., 2016; Srivatsan et al., 2019; Stirling et al., 2011; Yuen et al., 2007). We found relatively good correlations between Hug1 expression, Rnr3 expression and levels of Ddc2 foci, but poor correlation with the levels of Rad52 foci, consistent with the fact that Rad52 is not a member of the signaling pathway containing Ddc2, Rnr3, and Hug1. Mutations causing increased rates in GCR assays and other genome stability assays fell into two groups: those that did or did not result in increased DDR markers. These results indicate that increased DDR markers and increased DNA repair markers do not always reflect the increases in DNA damage or pathway alterations that result in increased genome instability.

Remarkably, the mutations identified in different DDR marker screens were only partially correlated, even when the markers belong to the same pathway. These differences likely correspond to the ability of different pathways to affect individual responses, such as transcriptional repression of the *HUG1* and *RNR3* genes, the rate at which the induced Hug1-EGFP and Rnr3-GFP proteins are degraded, and the rate at which Ddc2 foci intermediates turnover. In addition, these differences could also be due to the relative sensitivity of the assays and the possible accumulation of additional mutations or suppressors during the propagation of the libraries of mutant strains that were tested. In spite of these complications, approximately 70% of the mutations identified as inducing both Hug1 and Rnr3 expression also caused increased levels of Ddc2 foci (**Fig. 3C**); note that many of the mutations that induced Hug1 and Rnr3 expression but did not cause increased Ddc2 foci directly affected transcriptional regulation and do not directly affect the DDR. Moreover, the mutations identified in all three of the Hug1 and Rnr3 expression and Ddc2 foci assays affected genes involved in HR and DSB repair (*RAD52*, *RAD51*, *RAD54*, *RAD55*, *MRE11*, *SGS1*, *RMI1*, *TOP3*, *ESC2*, *SLX5*, *SLX8*), DNA replication (*MRC1*, *POL32*, *RAD27*), prevention of DNA replication errors and/or processing stalled replication forks (*DIA2*, *RAD5*, *RRM3*, *WSS1*, *MMS1*, *MMS22*, *RTT101*, *RTT107*), as well as affecting individual genes involved in processes that were not expected (*ELM1*, *IRA2*, *RTT109*, *SIC1*, *SPT21*). 24 mutations comprising the majority (89%) of mutations identified in all three of these DDR marker assays also caused increased GCR rates in at least one GCR assay, although mutations in some of these genes only cause very modest rate increases (*MRC1*, *RAD54*, *RRM3*, *POL32*).

In contrast, the mutations that induced the accumulation of Rad52 foci did not show a clear correlation with the mutations that induced the Hug1, Rnr3 and Ddc2 markers. This poor correlation is likely due to the fact that the accumulation Rad52 foci is not simply a measure of increased levels of DNA damage but rather measures damage that is being repaired by HR and the relative lifetime/turnover of these HR intermediates. None of these factors are unique to Rad52 foci and hence are likely to be also be shared with other DNA repair proteins that have been used as foci markers of the DNA damage response, including proteins like BRCA1, BRCA2, RAD51, MDC1, 53BP1 and RPA (Fernandez-Capetillo et al., 2002; Lou et al., 2003; Paull et al., 2000; Stewart et al., 2003; Wang et al., 2002; Zhong et al., 1999). In spite of these complications, a subset of the high confidence mutations that cause increased DNA damage signaling detected in the Hug1, Rnr3, and Ddc2 assays also cause increased levels of Rad52 foci, including mutations affecting *MMS1*, *MRE11*, *POL32*, *RAD27*, *RAD51*, *RAD54*, *RAD55*, *RMI1*, *RTT101*, *RTT107*, *RTT109*, *SGS1*, *SLX8*, and *WSS1*, all of which could alter the kinetics of DNA repair resulting in increased repair intermediates that both activate DNA damage signaling and are substrates for Rad52. Interestingly, the more DDR assays a mutation was identified in, the more likely the mutation was to caused increased GCR rates, with mutations identified in 4 or 5 DDR assays with one exception always causing increased GCR rates (**Suppl. Fig. 7**); this suggests that these mutations either cause high levels of DNA damage or cause a type of DNA damage that is particularly prone to mis-repair resulting in the formation of GCRs.

DDR markers as measured by cytological assays indicate the presence of DNA damage but cannot reveal if the damage is repaired in a conservative manner and hence does not result in increased genome instability or if it is repaired in a non-conservative manner leading to the formation of GCRs. Moreover, cytological assays measure the induction of DDR markers in the bulk of the cells (e.g. the “average” response), whereas genetic assays detect increased levels of “rare events”, which often not observable in the bulk of the cells. Thus, even in the case of mutations that give rise to both increased levels of DNA damage as measured cytologically and increased GCR rates, the GCRs measured at a per cell level are lower than the measured levels of cells exhibiting increased DNA damage markers. Hence, either most of the damage in these cells is either properly repaired and does not lead to the formation of GCRs or leads to the formation of GCRs such as reciprocal translocations that cannot be detected in the assays used. The fact that cells bearing these mutations are viable suggests that much of this damage is successfully repaired without formation of GCRs.

One of the reasons for performing the present study was to determine if assessing the effects of GCR-inducing mutations on the DDR could provide insights into the defects caused by these mutations that result in increased GCR rates in cases where the cause of increased GCR rates is not understood. Strikingly, the effects of the mutations tested in the DDR marker assays showed a limited correlation with their effects on the accumulation of GCRs. The mutations could be placed into four categories: GCR+ DDR+, GCR+ DDR-, GCR- DDR+, and GCR- DDR- (**Fig. 7**). Because the DDR+ category is based on identification of a mutation in one or more assays, the DDR+/DDR- status for mutations observed in less than two assays is sensitive to false positive and false negative errors in the individual screens. The GCR+ DDR+ mutations included most of the mutations in replication genes, genes encoding structure-specific endonucleases that act on normal and damaged replication intermediates, genes encoding the Smc5-Smc6 complex, and HR genes (**Suppl. Fig 7, Suppl. Table 5**). It seems likely that these mutations result in increased replication errors that underlie GCRs or increased steady state levels of damaged DNAs due to failure of correct repair that are then channeled into GCRs. The more surprising GCR+ DDR- mutations comprising 87 mutations could be explained if they increase the rate of aberrant repair acting on normal levels of endogenous DNA damage leading to increased formation of GCRs (**Suppl. Table 5**). Notable examples of such mutations include: 1) *pif1* that causes defects in suppression of *de novo* telomere addition reactions at DNA damage; 2) different checkpoint mutations that disrupt the DDR and cell cycle delay/arrest in response to DNA damage (*chk1*, *ddc2*, *dun1*, *mec1*, *mec3*, *mrc1-aq*) allowing aberrant repair to occur; 3) translesion synthesis, which may prevent the accumulation of DNA breaks during replication caused by an inability of the replicative DNA polymerases from extending past lesions or misincorporated bases (*rad30*, *rev1*, *rev3*, *rev7*, *srs2*); and, 4) mutations like *exo1* that by altering DSB resection but not altering the efficiency of DNA repair could change the balance between HR and end joining reactions that could lead to GCRs. There were a large number of GCR- DDR+ mutations, although many of these mutations were found in only one DDR assay and could represent false positives. Of the smaller number of mutations identified in multiple DDR assays that represent the best candidate mutations for only activating the DDR without increasing GCR rates, these could cause increased DNA damage that is not acted on by repair pathways that can produce GCRs or they could alter the kinetics of the DDR resulting in increased activation of the DDR without effecting the efficiency of repair itself. Examples of such mutations include: 1) those affecting the proteosome that could alter the turnover of DDR proteins; and, 2) mutations like *yku70* and *yku80* that could alter activation of the DDR and even reduce NHEJ that contributes to the formation of GCRs. Regardless, the number of GCR- DDR+ mutations identified indicate that activation of the DDR alone is not a reliable indicator of increased genome instability.

**Figure 7.**
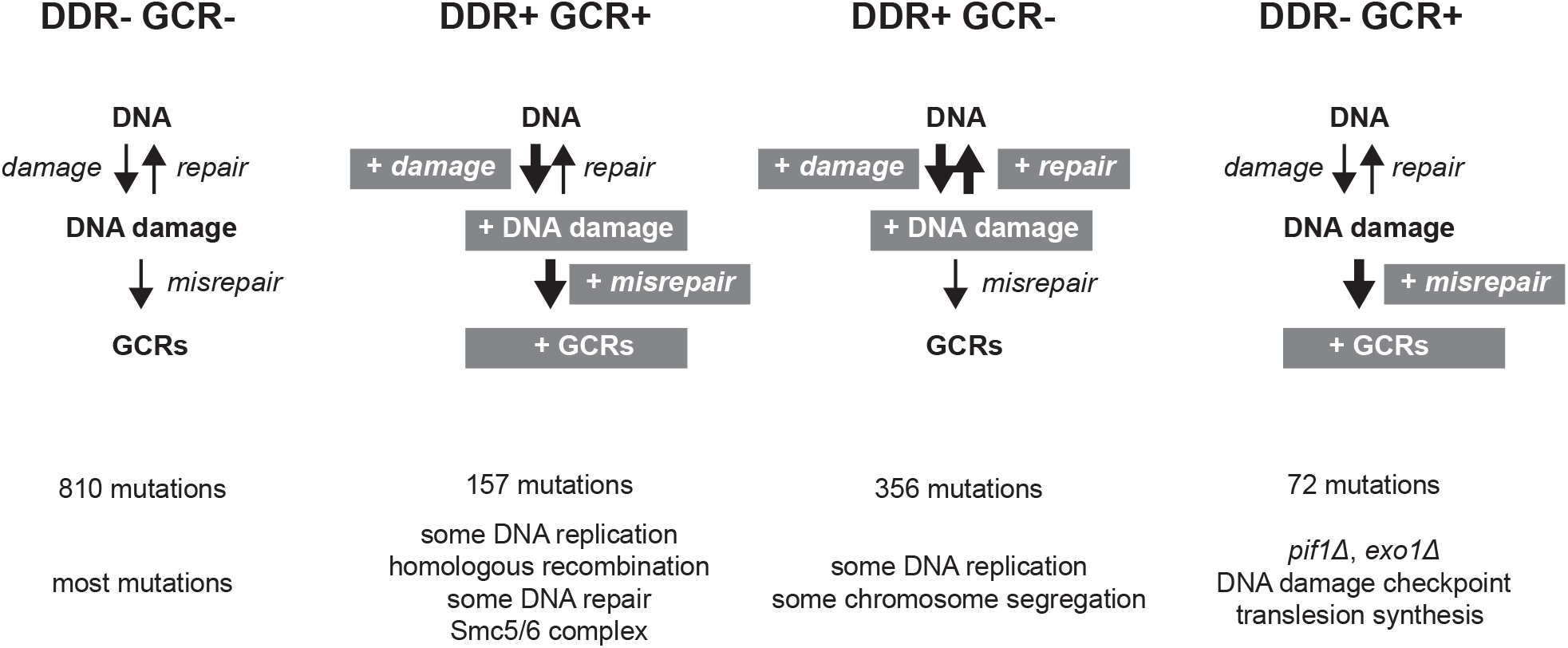
Classification of analyzed mutations. Mutations analyzed by at least one DDR assay and at least one GCR assay were categorized into DDR- GCR-, DDR+ GCR+, DDR+ GCR-, and DDR- GCR+ categories.

## AUTHOR CONTRIBUTIONS

Conceptualization, C.D.P. and R.D.K.; Methodology, B.Z.L., C.D.P. and R.D.K.; Investigation, B.Z.L. and C.D.P.; Formal Analysis C.D.P.; Data Curation C.D.P.; Writing – Original Draft, C.D.P.; Writing – Review & Editing, C.D.P. and R.D.K.; Funding Acquisition, R.D.K. and C.D.P.

## ACKNOWLEDGEMENTS

This work was funded by NIH R01 GM26017 to R.D.K. and the Ludwig Institute for Cancer Research to R.D.K. and C.D.P. The authors would like to thank Dr. John Petrini (Sloan Kettering Institute) for helpful comments on the manuscript.

## STAR METHODS

### KEY RESOURCES TABLE

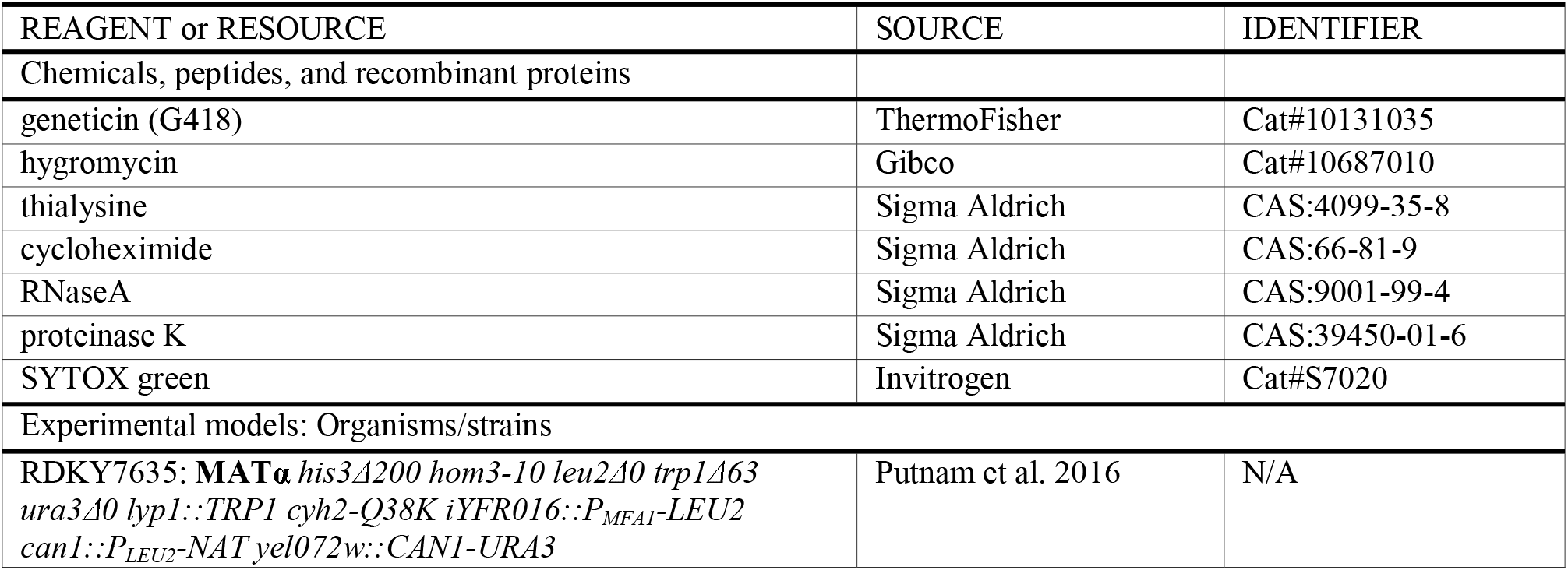
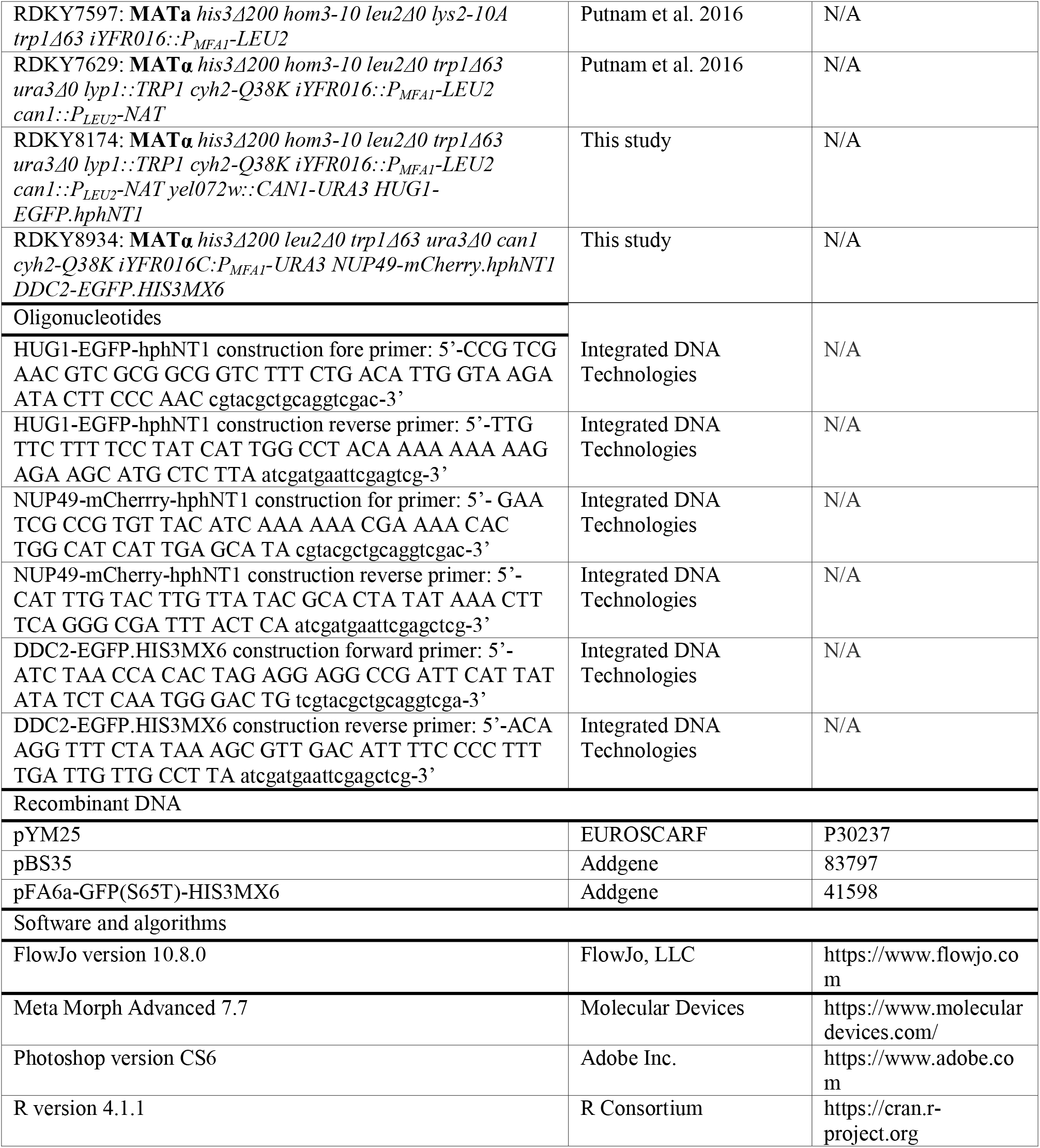

### RESOURCE AVAILABILITY

#### Lead contact

Further information and requests for resources and reagents should be directed to and will be fulfilled by the lead contact, Christopher Putnam.

#### Materials availability

Bait strains used in the systematic crosses are available from the lead contact upon request.

#### Data and code availability

Data are provided in the supplemental tables. Any additional information required to reanalyze the data reported in this paper is available from the lead contact upon request.

### METHOD DETAILS

#### Bait strain construction

The bait strain containing the *HUG1-EGFP* marker was created by integrating a *EGFP-hphNT1* cassette amplified by PCR from pYM25 (EUROSCARF P30237) using the primers 5’-CCG TCG AAC GTC GCG GCG GTC TTT CTG ACA TTG GTA AGA ATA CTT CCC AAC cgtacgctgcaggtcgac-3’ and 5’- TTG TTC TTT TCC TAT CAT TGG CCT ACA AAA AAA AAG AGA AGC ATG CTC TTA atcgatgaattcgagtcg-3’ onto the 3’ end of the *HUG1* coding sequencing in the strain RDKY7635 (Putnam et al., 2016) to create RDKY8174 (**MAT**α *his3*Δ*200 hom3- 10 leu2*Δ*0 trp1*Δ*63 ura3*Δ*0 lyp1::TRP1 cyh2-Q38K iYFR016::P_MFA1_-LEU2 can1::P_LEU2_-NAT yel072w::CAN1-URA3 HUG1-EGFP.hphNT1*). The bait strain containing the *DDC2-EGFP* marker was created by: [1] crossing RDKY7597 with RDKY7629 (Putnam et al., 2016), [2] replacing the *iYFL016C::P_MFA1_-LEU2* with a *iYFL016C::P_MFA1_-URA3* in a **MATa** haploid spore clone, [3] re-crossing the resulting strain to a **MAT**α haploid spore clone from the same RDKY7597 x RDKY7629 cross, [4] screening for a **MAT**α haploid spore clone with the *iYFL016C::P_MFA1_-URA3* marker, [5] selecting for a mutation in *CAN1* by plating on CSM-Arg plates containing 60 μg/mL canavanine (Sigma Aldrich), [6] integrating a PCR product containing the *mCherry-hphNT1* fragment from pBS35 (Addgene #83797) with the primers 5’- GAA TCG CCG TGT TAC ATC AAA AAA CGA AAA CAC TGG CAT CAT TGA GCA TA cgtacgctgcaggtcgac-3’ and 5’- CAT TTG TAC TTG TTA TAC GCA CTA TAT AAA CTT TCA GGG CGA TTT ACT CA atcgatgaattcgagctcg-3’, and [7] integrating a PCR product containing the *EGFP-HIS3* fragment amplified from pFA6a-GFP(S65T)-HIS3MX6 (Addgene #41598) with the primers 5’- ATC TAA CCA CAC TAG AGG AGG CCG ATT CAT TAT ATA TCT CAA TGG GAC TG tcgtacgctgcaggtcga-3’ and 5’-ACA AGG TTT CTA TAA AGC GTT GAC ATT TTC CCC TTT TGA TTG TTG CCT TA atcgatgaattcgagctcg-3’. The resulting strain was RDKY8934 (**MAT**α *his3*Δ*200 leu2*Δ*0 trp1*Δ*63 ura3*Δ*0 can1 cyh2-Q38K iYFR016C:P_MFA1_-URA3 NUP49-mCherry.hphNT1 DDC2-EGFP.HIS3MX6*).

#### Systematic genetic crosses

The *HUG1-EGFP* containing bait strain was crossed twice to a selected set of the *S. cerevisiae* deletion collection (BY4741 strains, **MATa** *his3*Δ*1 leu2*Δ*0 met15*Δ*0 ura3*Δ*0*) and once to the entire collection of BY4741 **MATa** strains using a RoToR pinning robot (Singer Instruments). The *DDC2-EGFP.HIS3MX6* containing bait strain was crossed to a set of 468 BY4741 **MATa** deletion strains containing mutations selected as causing increased Hug1-EGFP expression or increased GCR rates or the *leu2*Δ*::kanMX4* control mutation. For crossing the *HUG1-EGFP* containing bait strain to the deletion collection, the bait strain was grown on YPD and crossed in quadruplicate with arrayed deletion strains by pinning onto fresh YPD agar plates using a Singer RoToR followed by growth for two days. Cells were subjected to two rounds of pinning onto diploid selection medium (YPD-agar containing 200 μg/mL geneticin (G418, ThermoFisher) and 200 μg/mL hygromycin (Gibco)) and grown for 1-2 days at 30°C. The cells were then pinned onto pre-sporulation medium (15 g Difco nutrient broth (BD Biosciences), 5 g Bacto-yeast extract (ThermoFisher), 10 g Bacto-agar (ThermoFisher), and 62.5 mL 40% glucose (ThermoFisher) per 500 mL) and grown for 3 days at 30°C. Cells from the pre-sporulation medium were then pinned onto sporulation medium (10 g potassium acetate, 0.05 g zinc acetate, 20 g Bacto-agar per liter, containing a final concentration of 50 μg/mL hygromycin) and incubated for 7 days at 30°C. The resulting spore-containing cells were then subjected to two rounds of pinning onto diploid killing medium (1.7 g yeast nitrogen base without amino acids and without ammonium sulfate (BD Biosciences), 1 g L-glutamic acid monosodium salt, 2 g CSM dropout mix without lysine and leucine (US Biological), 20 g Bacto-agar, 50 mL of 40% glucose per liter, containing a final concentration of 50 μg/mL thialysine (Sigma Aldrich), 10 μg/mL G418, and 200 μg/mL hygromycin) followed by growth for 5 days at 30°C for the first pinning and 2 days at 30°C for the second pinning. Cells were then subjected to two rounds of pinning and growth on haploid selection medium (1.7 g yeast nitrogen base without amino acids and without ammonium sulfate, 1 g L-glutamic acid monosodium salt, 2 g CSM dropout mix without leucine, 20 g Bacto-agar, 50 mL of 40% glucose per liter, containing a final concentration of 200 μg/mL G418 and 200 μg/mL hygromycin) and grown for 2 days at 30°C. Then the cells were pinned and grown on YPD-agar followed by storage at −-80°C in YPD media containing 20% (v/v) glycerol. Crossing the *DDC2-EGFP.HIS3MX6* containing bait strain was performed by essentially the same procedure, except for the composition of the diploid killing medium (1.7 g yeast nitrogen base without amino acids and without ammonium sulfate, 1 g L-glutamic acid monosodium salt, 2 g CSM dropout mix without uracil and histidine, 20 g Bacto-agar, 50 mL of 40% glucose per liter, containing a final concentration of 50 μg/mL canavanine, 10 μg/mL cycloheximide, 200 μg/mL G418, and 200 μg/mL hygromycin) and the haploid selection medium (1.7 g yeast nitrogen base without amino acids and without ammonium sulfate, 1 g L-glutamic acid monosodium salt, 2 g CSM dropout mix without uracil and histidine, 20 g Bacto-agar, 50 mL of 40% glucose per liter, containing a final concentration of 200 μg/mL G418 and 200 μg/mL hygromycin).

#### DNA content analysis

Cell-cycle analysis was conducted as previously described (Vanoli et al., 2010). In brief, 1×10^7^ cells were collected by centrifugation and resuspended in 70% ethanol for 16 h. Cells were then washed in 0.25 M Tris-HCl (pH 7.5), resuspended in the same buffer containing 2 mg/ml of RNaseA (Sigma Aldrich) and incubated at 37°C for at least 1 h, then treated overnight with proteinase K (1 mg/ml; Sigma Aldrich) at 37°C. Cells were collected by centrifugation, resuspended in 200 mM Tris-HCl (pH 7.5) buffer containing 200 mM NaCl and 80 mM MgCl_2_ and stained in the same buffer containing 1 μM SYTOX green (Invitrogen). Samples were then diluted 10-fold in 50 mM Tris-HCl (pH 7.8) and analyzed using a Becton Dickinson FACScan instrument. This FACS analysis verified that all of the strains used in the experiments reported in this study were haploids.

#### Measurement of Hug-EGFP levels

To measure Hug1-GFP expression, logarithmic-phase cultures grown in YPD medium in 96 well plates, centrifuged, and the cells were resuspended in 100 μL sterile water and sonicated in a Cole-Palmer 8891 Water Bath Sonicator. The cells were then analyzed using a BD LSR Fortessa analytical cytometer with a High-Throughput Sampler. Excitation was at 488 nm, and the fluorescence signal was collected through a 505-nm long-pass filter and a HQ510/20 band-pass filter (Chroma Technology Corp). For each sample, 30,000 events were recorded. The mean value of GFP abundance was calculated using FlowJo software and normalized to the mean GFP value of wild-type cells.

#### Measurement of Ddc2-GFP foci

Cells were grown in complete synthetic medium (Sigma Aldrich) to log phase and examined by live imaging using an Olympus BX43 fluorescence microscope with a 60x 1.42 PlanApo N Olympus oil-immersion objective. GFP fluorescence was detected using a Chroma FITC filter set and captured with a Qimaging QIClick CCD camera. Images were analyzed using Meta Morph Advanced 7.7 imaging software, keeping processing parameters constant within each experiment.

#### Cell cycle distribution data analysis

##### Data compilation

The cell cycle distribution from log phase cells was obtained for the **MATa** BY4741 haploid deletion collection and the homozygous diploid BY4743 deletion collection from previously published data (Hoose et al., 2012; Koren et al., 2010; Soifer and Barkai, 2014). For the diploid deletion collection data, the published supplement containing cell cycle parameters from the Dean-Jett-Fox model (Fox, 1980) as implemented in FlowJo were used (Hoose et al., 2012). For the haploid deletion collection data, FACS data were downloaded from the FlowRepository (Spidlen et al., 2012) and were analyzed using the Denn-Jett-Fox model in FlowJo version 10 (FlowJo, 2019) after removing doublets that ran perpendicular to the laser by gating on a forward scatter height (FSC-H) vs. forward scatter width (FSC-A) plot. Note that cell-cell doublets that are parallel to the laser cannot be excluded by this gate. Cell cycle distribution data were filtered for problematic observations. These observations included those flagged by the original authors (Hoose et al., 2012), those with problematic fits based on visual inspection, those with very small numbers of cells, or those with abnormal ploidies. All observations, including those filtered out, are reported in **Supplemental Table 4**.

##### Determination of aggregation-corrected cell-cycle distributions

Ternary plots are ideal ways to analyze data in which three variables sum to a constant; in the case of cell cycle data, we can scale the percent of G1, S, and G2 cells so that they sum to a constant 100%. This scaling can be performed by ignoring the sub-G1 and super-G2 populations and dividing the percent of G1, S, and G2 cells by their sum. This scaling, however, ignores the information in the super-G2 population about cell-cell aggregation present in the sample, which would tend to systematically decrease the overall levels of the G1 and S populations, as the aggregates with the smallest DNA content, two G1 cells, will be scored with the G2 population, and aggregates with increased DNA content will be scored with the super-G2 population. To take aggregation into account, distributions with a super-G2 population were fitted with by a model in which cell-cell aggregates were assumed to be cell-cycle independent and aggregates larger than two cells were ignored, as they were assumed to be present at vanishingly small amounts in the sample. In this model, the underlying fractions of cells in the G1, S, and G2 populations (*f_G1_*, *f_S_*, *f_G2_*, where *f_G1_*+*f_S_*+*f_G2_*=1) and the fraction of cells in aggregates (0≤*f_A_*≤1) were determined by fitting the following equations:

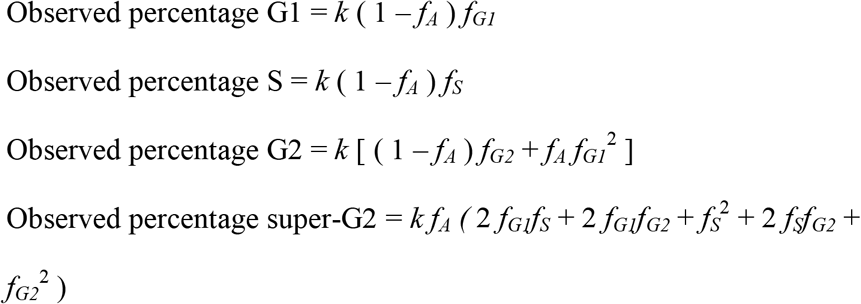

where *k* is the sum of the observed percentages of G1, S, G2, and super-G2 cells. These four equations have only three unknowns (*f_G1_*, *f_S_*, *f_A_*), and the parameters were determined using Powell minimization as implemented in the SciPy python package (Virtanen et al., 2020). In general, the fitted values of *f_G1_*, *f_S_*, and *f_G2_* were only modestly different from those calculated by scaling the sum of the G1, S, and G2 populations to 1, even in the presence of moderate fractions of aggregation. Downstream analyses used the aggregation-corrected frequencies *f_G1_*, *f_S_*, and *f_G2_*.

##### Projecting cell cycle data onto a ternary plot

With each observation expressed as the fractional distribution of cells in each cell cycle phase (*f_G1_*, *f_S_*, *f_G2_*, where *f_G1_*+*f_S_*+*f_G2_*=1), each observation was represented as a two-dimensional point on a ternary plot. With G1 at top vertex of the graph (0.5, (3^1/2^)/2), S at left vertex (0, 0), and G2 at the right vertex (0, 1), the fractional distribution can be related with a point (*x*, *y*) on the ternary plot using the following equations:

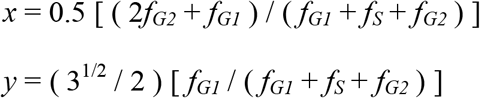

##### Merging multiple observations

To merge multiple observations together, the geometric median of all projected observations in the ternary plot was determined using Weiszfeld’s algorithm, which iteratively re-weights points to minimize the weighted sum of squares of the distances of the points with the median (Weiszfeld, 1937). This merging procedure was performed on multiple observations of an allele, but in principle could be performed on all observations of all alleles affecting a gene, a complex, or a pathway. The resulting median in the ternary plot (*x_m_*, *y_m_*) was converted back to a fractional distribution (*f_G1_*, *f_S_*, *f_G2_*) using the relationships:

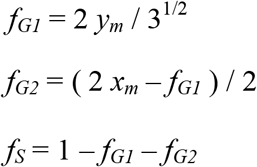

##### Identification of mutations causing altered cell-cycle distributions

First, a null model was generated using the ternary plot distances between the geometric median of the control strain observations and each individual observation. These distances had a non-normal distribution, with reduced observations at low distances and a distribution skewed to longer distances (**Suppl. Fig. 2B**). Because these distances can be modeled as arising from deviations in two dimensions, we compared the control distances to a theoretical distribution of distances arising from points in which the *x* and *y* values were normally distributed about the origin with a standard deviation of 1 (**Suppl. Fig 2B**). A linear relationship in the quantile-quantile plot at short quantiles suggests that the observed distributions fit this theoretical model with a subset of outliers at large values. From the quantile-quantile plot, the standard deviation for the control distances in haploid strains was 0.0263 and the diploid strains was 0.0340 (**Suppl. Fig 2C**).

Next, all of the observations for each deletion allele were compared to the null models. Haploid strain observations were compared to the haploid null model, and diploid strain observations were compared to the diploid null model. For a deletion allele with *n* observations, each the deviation vector for each observation *i,* Δ***p_i_***, was calculated:

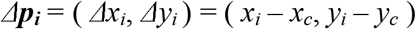

where *x_i_* and *y_i_* are the ternary plot *x* and *y* values for the observation and *x_c_* and *y_c_* correspond to the *x* and *y* values of the geometric median of the control observations. All of the observed deviation vectors for the deletion allele were then summed, and the length of the summed vector, ashok ***p***ashok was determined:

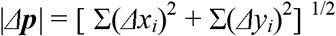

The observed length was then compared to a million simulations in which *n* deviation vectors were randomly generated, and the length of the sum of these randomly generated vectors was determined. For the generation of each simulated observation, the length of the vector was taken from the null model and the orientation was randomly assigned. The *p*-value for the change in the cell-cycle distribution was taken as the fraction of the million simulations that had summed deviation vectors with longer lengths than the observed length of the summed deviation vectors. The Benjamini-Hochberg false-discovery rate correction for multiple tests was then applied to the resulting *p*-values for all of the deletion alleles (Benjamini and Hochberg, 1995).

#### Quantification and statistical analysis

All statistical analyses were performed with R version 4.1.1. Significance for the Hug1-EGFP expression data was performed by fitting to a Gaussian distribution as described in the main text, Figure 1, and Supplemental Figure 1. Significance for the Ddc2-GFP foci data was performed using 95% cutoffs of the wild-type data as described in the main text and Figure 3. Significance for the cell cycle distribution was performed as described in the main text and methods based on the two-independent Gaussian error model.

