## Supplemental Figures and Text for "Identification of different classes of genome instability suppressor genes through analysis of DNA damage response markers"

By

Bin-Zhong Li[1], Richard D. Kolodner[1, 2, 3, 4] and Christopher D. Putnam[1, 5]

From

Ludwig Institute for Cancer Research[1], Department of Cellular and Molecular Medicine[2], Moores-UCSD Cancer Center[3], Institute of Genomic Medicine[4], and Department of Medicine[5], University of California San Diego School of Medicine, 9500 Gilman Drive, La Jolla, CA 92093-0669

Address correspondence to:
Christopher D. Putnam

(858) 534-5125 (phone)
(858) 534-7750 (fax)

### SUPPLEMENTAL FIGURE LEGENDS

**Supplemental Figure 1; related to Figure 1. Selection and analysis of Hug1-EGFP DDR marker expression in mutant strains. A.** Log-phase cells containing either a *DSE2-EGFP*, *RNR3-EGFP*, *DDR2-EGFP*, or a *HUG1-EGFP* marker were grown in the absence or presence of 100 mM HU to induce the DDR response. EGFP levels were then measured by FACS. Both *RNR3-EGFP* and *HUG1-EGFP* showed a substantial increase in expression levels after HU treatment. **B.** Independent Hug1-EGFP expression measurements are highly correlated. Subsets of the deletion collection were independently crossed with the *HUG1-EGFP* query strain and Hug1-EGFP expression in haploid progeny was measured by FACS. The data from three independent experiments was analyzed: "High Priority 1" (412 valid measurements), "High Priority 2" (482 valid measurements), and "Whole Genome" (4,936 valid measurements). Plots of the fold increase in Hug1-GFP expression for strains with the same genotypes from each experiment show that the observed values were highly correlated; Pearson correlation coefficient (*r*) and *p*-value for the null hypothesis that the true correlation is zero are shown for each pairwise comparison. **C.** Fold increases in Hug1-EGFP expression levels are normally distributed as shown by the linear relationship in the quantile-quantile plot that compares the quantiles of the experimental fold increase in Hug1-EGFP expression levels against the quantiles of a theoretical normal distribution. Outliers at higher quantiles correspond to strains with high levels of Hug1-EGFP expression. Regression of linear portion of the plot indicates that a fitted normal distribution of the data has a mean fold Hug1-EGFP of 1.05 and a deviation of 0.13. **D.** Histogram of all measured Hug1-EGFP expression levels relative to the control strain (open circles) fitted to a Gaussian distribution (red curve) described by the parameters from the quantile-quantile plot (panel C).

**Supplemental Figure 2; related to Figure 5. Ternary plot distance measurements identify mutants with altered cell cycle distributions. A.** Ternary plot diagrams of all haploid and diploid control strain observations; 292 and 219 observations are plotted for the haploid and diploid controls, respectively. **B.** (Left) Histogram of the theoretical distance distribution between the origin and two-dimensional points selected such that the x and y coordinates are selected from a normal distribution about 0 with a sigma of 1. (Middle and Right) Histograms of the distances in the ternary plots between the geometric median of the control strain observations and all individual control strain observations for haploid and diploid strains. **C.** A linear relationship in the quantile-quantile (q-q) plot demonstrates that the experimental distribution of distances of the control strain observations from the geometric median of these observations in the ternary plot matches the theoretical error model. The slope of the lines is the sigma value for the fitted distribution. **D.** Ternary plot of the median cell cycle distribution for each haploid mutant strain. **E.** (Top) Rank ordered distances between all haploid mutant strains and the geometric median of the control strain observations. (Bottom) The FDR values for the haploid mutant cell cycle distributions being different than the control strain distribution, which takes into account both changes in cell cycle distribution and the number of observations for that mutant (see Methods) **F.** Ternary plot of the median cell cycle distributions for each diploid mutant strain displayed as in panel B. **G.** Rank ordered distances and FDR values for all diploid mutant strains displayed as in panel E.

**Supplemental Figure 3; related to Figure 5. Cell cycle distributions of haploid DDR+ mutants. A,C,E,G,I.** Ternary plot diagram for the cell cycle distribution of haploid strains (Koren et al., 2010; Soifer and Barkai, 2014) colored by the effect of the mutation in these strains on induction of Hug1 (panel A), Rnr3 (panel C), Ddc2 foci (panel E), Rad52 foci in diploids (panel G), and Rad52 foci in haploids (panel I) (blue=no induction, red=induction). The black cross indicates the position of the wild-type control strain. Strains indicated outside the black circle have a significantly altered cell cycle distribution ($p<0.01$) compared to the wild-type control strain. **B,D,F,H,J.** Analysis of mutations causing induction of Hug1 (panel B), Rnr3 (panel D), Ddc2 (panel F), Rad52 foci in diploids (panel H), and Rad52 foci in haploid (panel J) based on the ternary plot distance between the mutant distribution and the geometric median of the control strain distribution. (Top) The cumulative distribution of all mutants (blue) is compared the cumulative distribution of distances for those mutations with significant induction of the relevant DDR marker. (Bottom) Plot of the fold DDR marker induction against the ternary plot distance. The vertical line indicates the significance cutoff for the cell cycle distribution.

**Supplemental Figure 4; related to Figure 5. Cell cycle distributions of homozygous diploid DDR+ mutants.** Cell cycle distributions of homozygous diploids (Hoose et al., 2012) associated with induction of the DDR markers Hug1 (**panels A,B**), Rnr3 (**panels C,D**), Ddc2 foci (**panels E,F**), Rad52 foci in diploids (**panels G,H**), and Rad52 foci in haploids (**panels I,J**). Figures are displayed as in Supplemental Figure 3.

**Supplemental Figure 5; related to Figure 5. Comparison of ternary plot distances caused by individual mutations with the number of DDR screens the mutations scored in.**

Mutations are plotted by the number of DDR screens they were identified in (x-axis) against the ternary plot distance (y-axis) for haploid (**panel A**) and diploid (**panel B**) data. Black dots are mutations that have been measured in at least one DDR assay. The solid line (y=0) corresponds to no distance between the mutation and the control in the ternary plot, and the dashed line is the threshold for significance (p=0.01). Red bars are the median GCR fold increase for mutations in each column.

**Supplemental Figure 6; related to Figure 6. Distribution of GCR rates and patch scores.** Distribution of the GCR rates and patch scores grouped by the effect of the mutations on Hug1 and Rnr3 expression; "+" indicates a significant effect in that assay, "-" indicates not significant, and "?" indicates not measured. The fraction above each category indicates the number of mutations causing a significant increase in genome instability as measured by GCR assays over the total number of mutations in that category.

**Supplemental Figure 7; related to Figure 6. Comparison of the fold GCR rate increase caused by individual mutations with the number of DDR screens the mutations scored in.** Mutations are plotted by the number of DDR screens they were identified in (x-axis) against the fold GCR rate increase (y-axis). Black dots are mutations that have been measured in at least one DDR assay; only some of these mutations are labeled, see Supplementary Table 5 for full mutation list. Grey dots are mutations that have not been measured in any DDR assay. The solid line (y=1) corresponds to no GCR fold increase, and the dashed line (y=3) is the threshold for significance. Red bars are the median GCR fold increase for mutations in each column. Note that the median is greater than the threshold for mutations identified in 3-5 DDR screens and that the median is near y=1 for mutations identified in 0-2 DDR screens. In general, the more DDR screens a mutation is identified in, the more likely the GCR fold increase will be significant, and almost all mutations identified in 4-5 DDR screens cause increased GCR rates. In spite of this, there are a number of GCR+ DDR- mutations (black dots in the number of significant DDR assays = 0 column) and the GCR fold increase caused by many of these mutations is as high or higher than that caused by many of the GCR+ DDR+ mutations; see Supplementary Table 5 and main text for details.

**Supplemental Table 1. Hug1-GFP observations for mutant strains.**

**Supplemental Table 2. Comparison of Hug1 and Rnr3 assays.**

**Supplemental Table 3. Ddc2-GFP foci observations for mutant strains.**

**Supplemental Table 4. Cell cycle distribution observations for mutant strains.**

**Supplemental Table 5. Merged mutant strain information.**

### SUPPLEMENTAL APPENDIX

**Analysis of cell cycle distribution alterations.** Cell cycle distribution data of unperturbed log-phase cells was derived from published FACS analysis of the *S. cerevisiae* haploid (Koren et al., 2010; Soifer and Barkai, 2014) and diploid (Hoose et al., 2012) deletion collections. Cell cycle distributions were taken from the published values for the diploid dataset (5,148 observations of 3,816 strains) or redetermined from the raw FACS data for the haploid dataset (7,327 observations of 4,698 strains; **Suppl. Table 4**; **see Methods**). Each cell cycle distribution (**Fig. 5A**) was fitted so that the percentage of cells in the G1-, S-, and G2-phases of the cell cycle summed to 100% after taking into account cell-cell aggregation (**Fig. 5B; see Methods**). Because of the constraint that $f_{G1}$+$f_S$+$f_{G2}$=1, the fitted distribution could be plotted as single points on a ternary plot diagram (**Fig. 5C; see Methods**). The ternary plot allowed for a direct method of merging multiple observations while retaining the $f_{G1}$+$f_S$+$f_{G2}$=1 constraint by determining the geometric median of the points for any one mutation (**see Methods**). Using the geometric median, the percentage of G1-, S-, and G2-phase cells in control measurements was found to be 21.5%, 25.1%, and 53.4% for the haploid data (n=292), and 22.8%, 24.3%, and 52.3% for the diploid data (n=219, **Suppl. Fig. 2A**).

The deviations of the individual control measurements from the geometric mean in the ternary plot fit an error model in which the deviations in the x and y directions are modeled as two independent Gaussian distributions (**Suppl. Fig. 2B**). The deviations of the control observations from their geometric median fit this distribution as observed by a similar distance histograms and linear quantile-quantile plot (**Suppl. Fig. 2B,C**). Using the fitted error model, p-values were assigned to individual observations in both the haploid and diploid datasets. This analysis identified 698 mutations in the haploid dataset and 350 mutations in the diploid dataset with an FDR of less than 0.01 and a ternary plot distance that was 3 or more times the standard deviation, which corresponds to $p<0.01$ in the two independent Gaussian error model. Increased fractions of G1 phase cells were caused by many defects affecting translation, including mutations in genes encoding ribosomal subunits, ribosomal RNA transcription, and ribosome maturation (**Suppl. Table 4)**, which is consistent with previous observations (Hoose et al., 2012; Moore, 1988; Popolo et al., 1982; Soifer and Barkai, 2014). In addition, mutations affecting mitochondrial function, adenine biosynthesis, NAD biosynthesis, and vacuole acidification also resulted in increased fractions of G1 phase cells (**Suppl. Table 4**). Increased fractions of S phase cells were observed for mutations affecting cell cycle progression, nucleotide biosynthesis, DNA replication, and histone synthesis and assembly (**Suppl. Table 4**). Increased fractions of G2 phase cells were observed for mutations affecting several DNA repair pathways, including HR, post-replication repair, the Mre11-Rad50-Xrs2 complex, the Sgs1-Top3-Rmi1 complex, and the Mms1-Mms22-Rtt101-Rtt107 cullin complex (**Suppl. Table 4**). In addition to DNA repair defects, defects in pathways involved in mitosis also increased the fraction of G2 phase cells, including mutations affecting cell cycle progression, chromosome cohesion and segregation, the actin cytoskeleton, karogamy, nuclear migration, and cytokinesis (**Suppl. Table 4**).

Supplemental Figure 1.

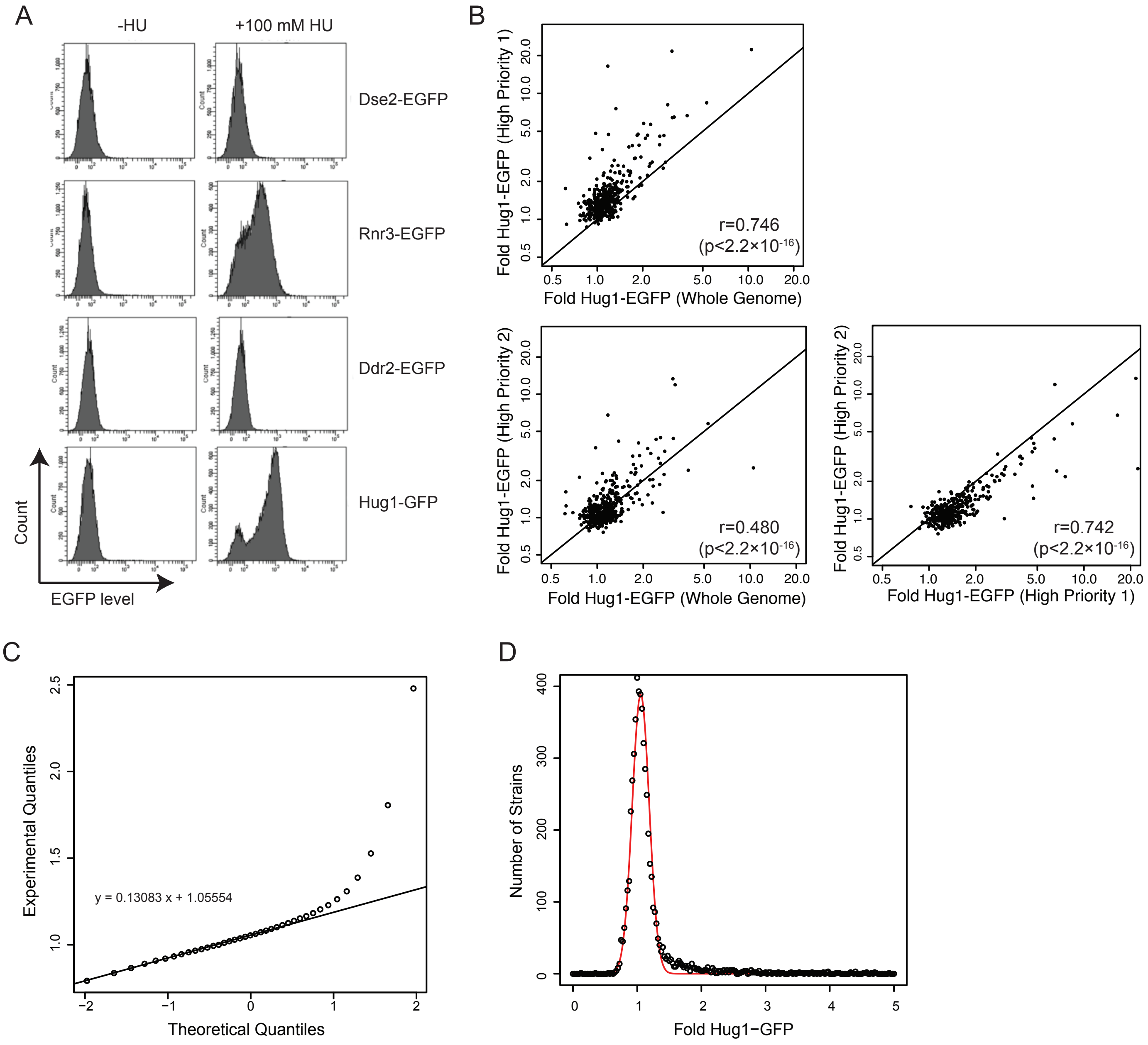

Supplemental Figure 2

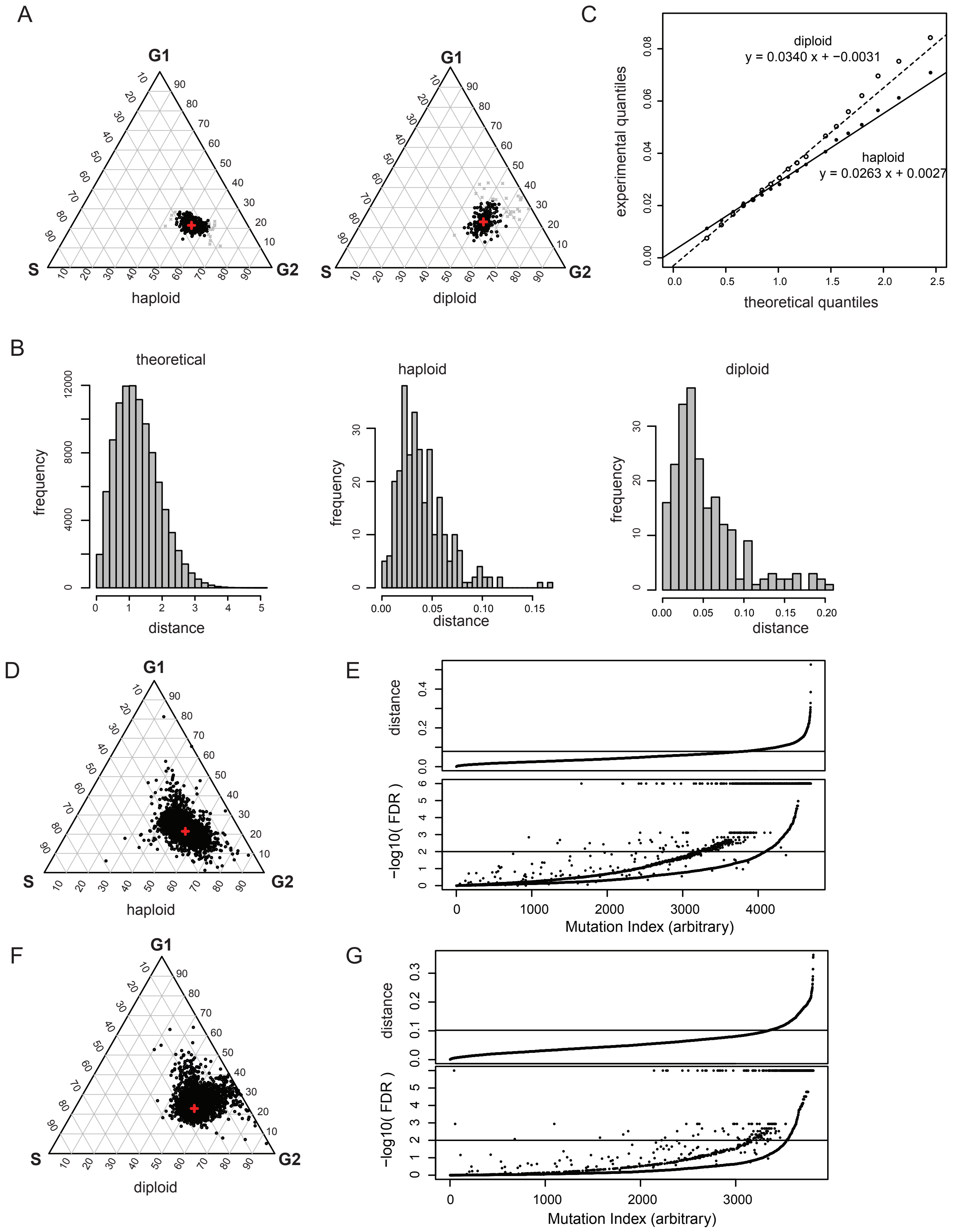

Supplemental Figure 3

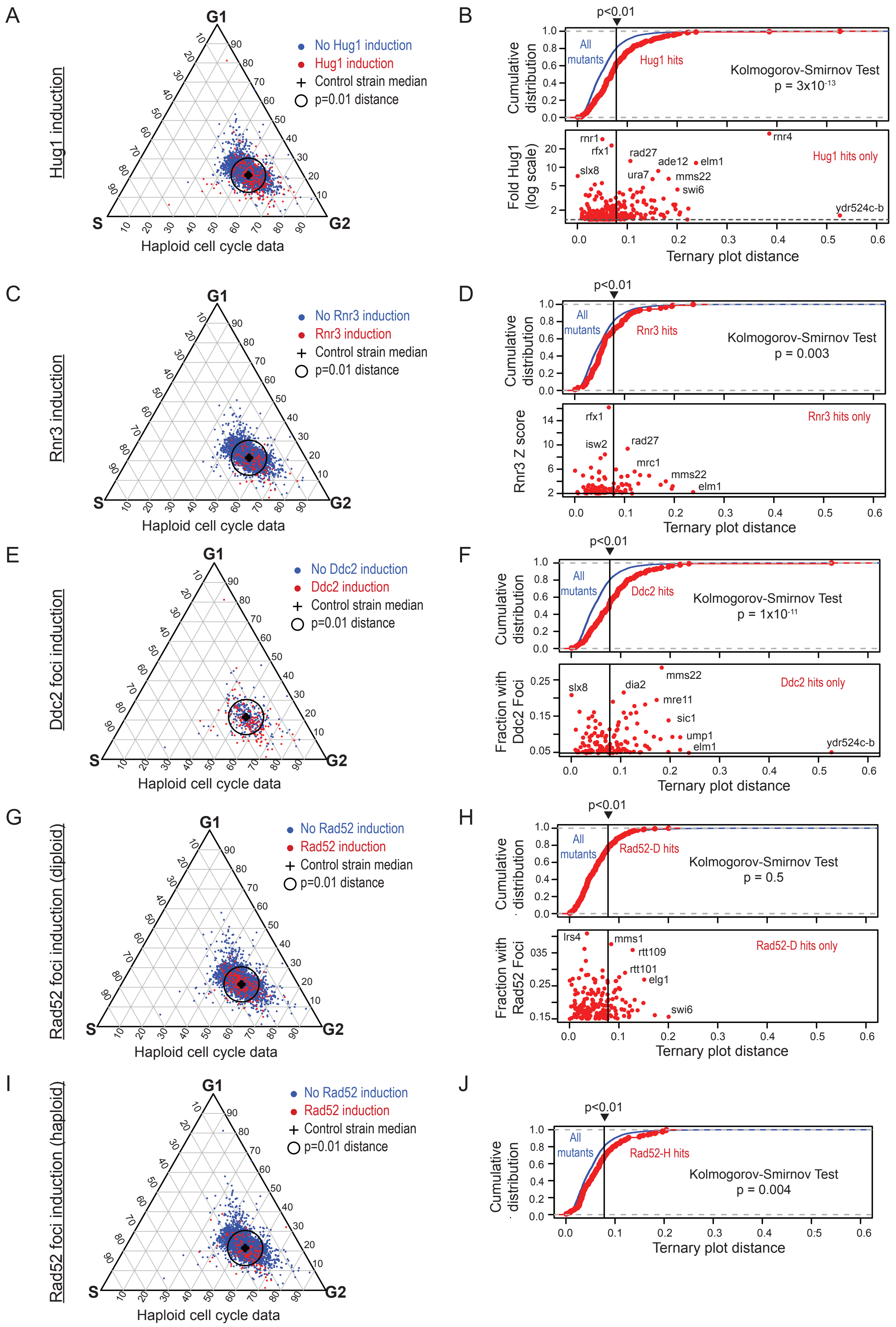

Supplemental Figure 4

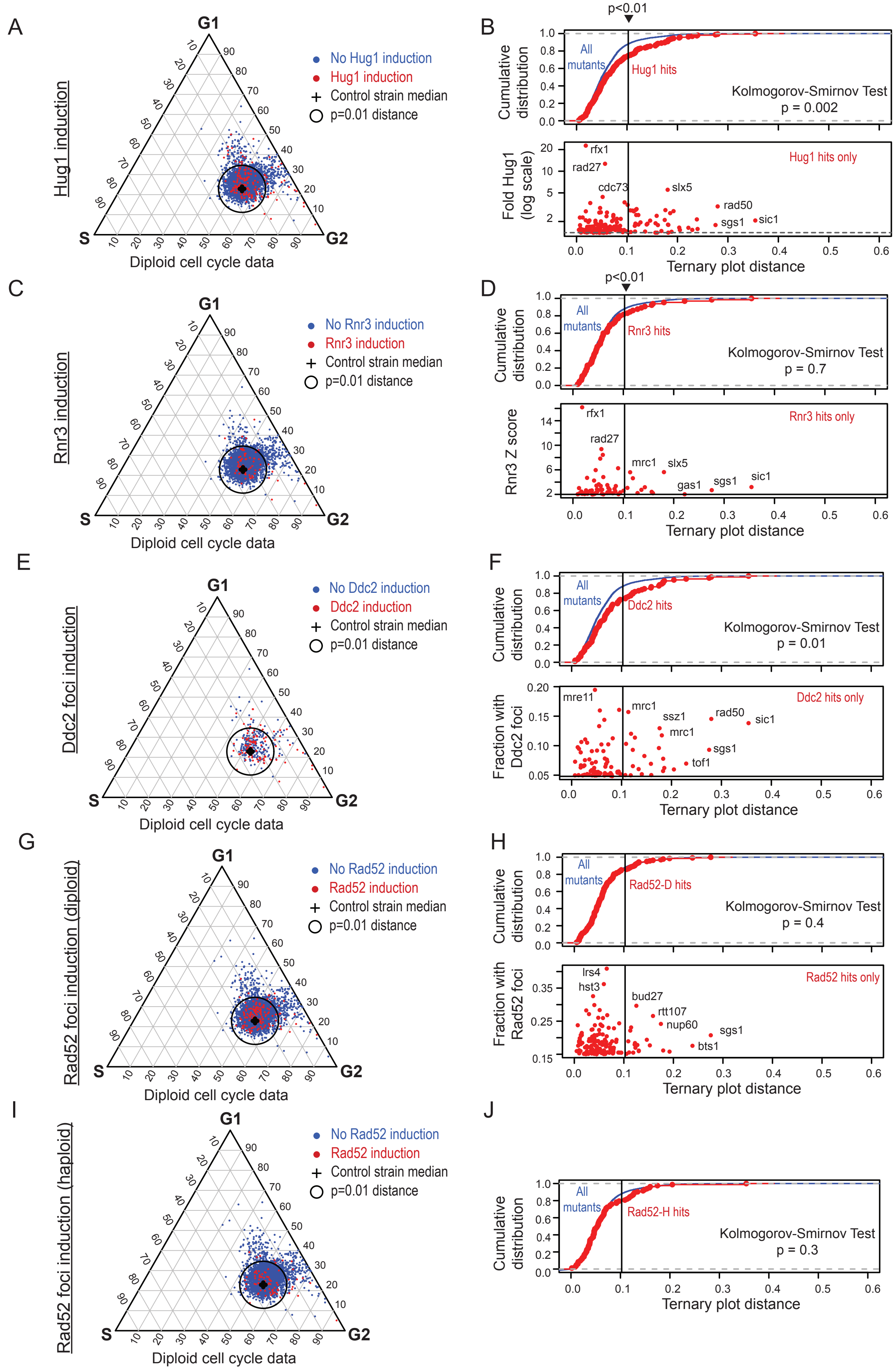

Supplemental Figure 5

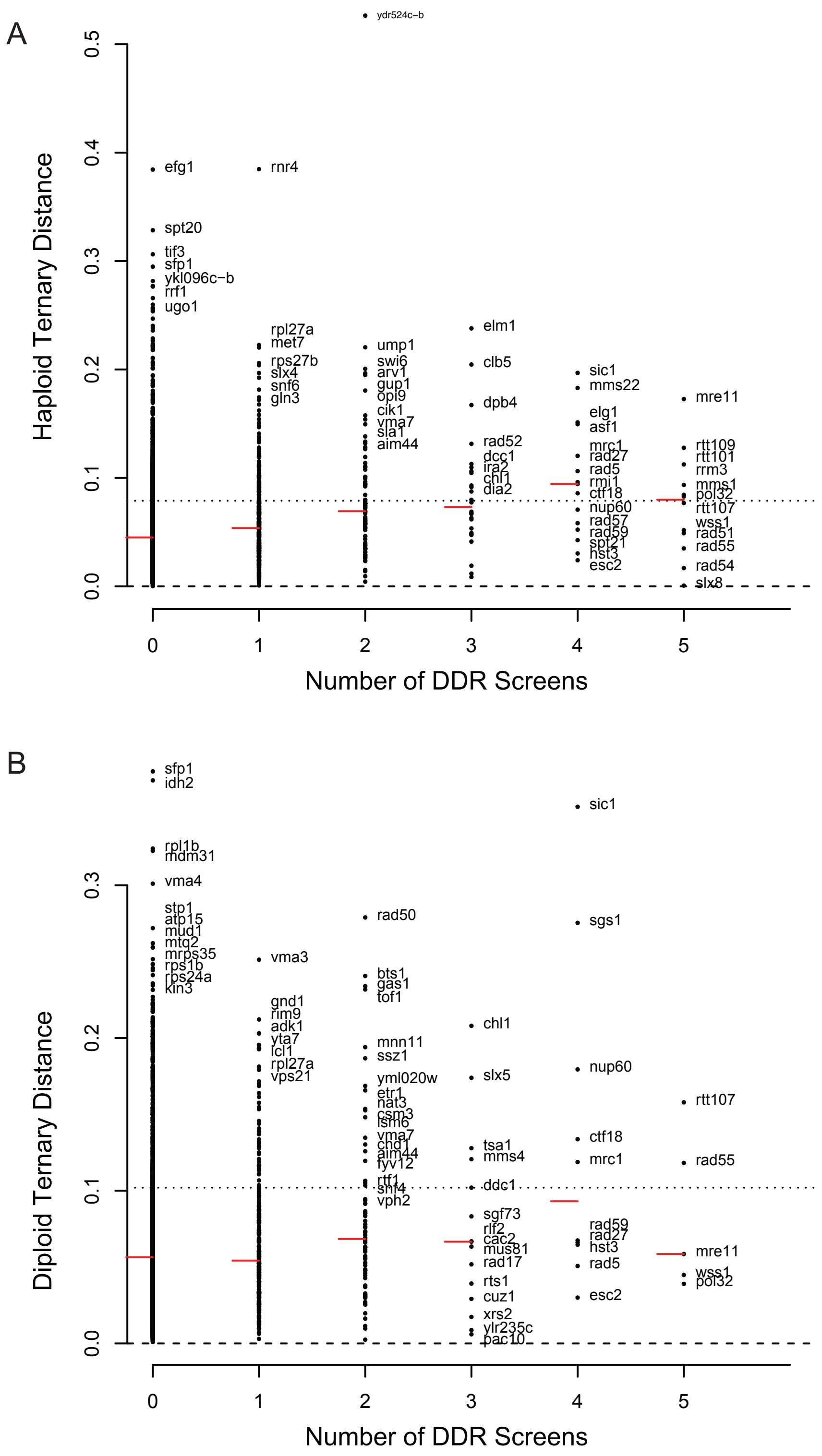

Supplemental Figure 6

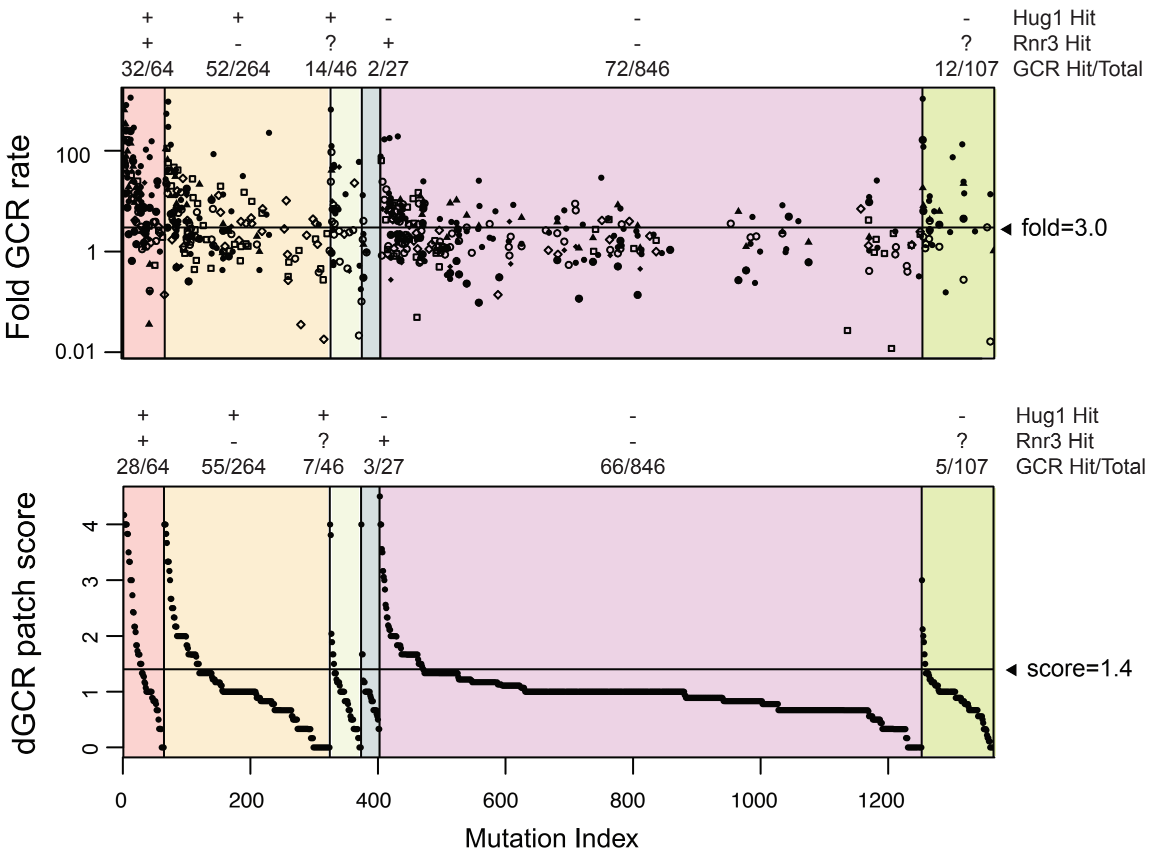

Supplemental Figure 7

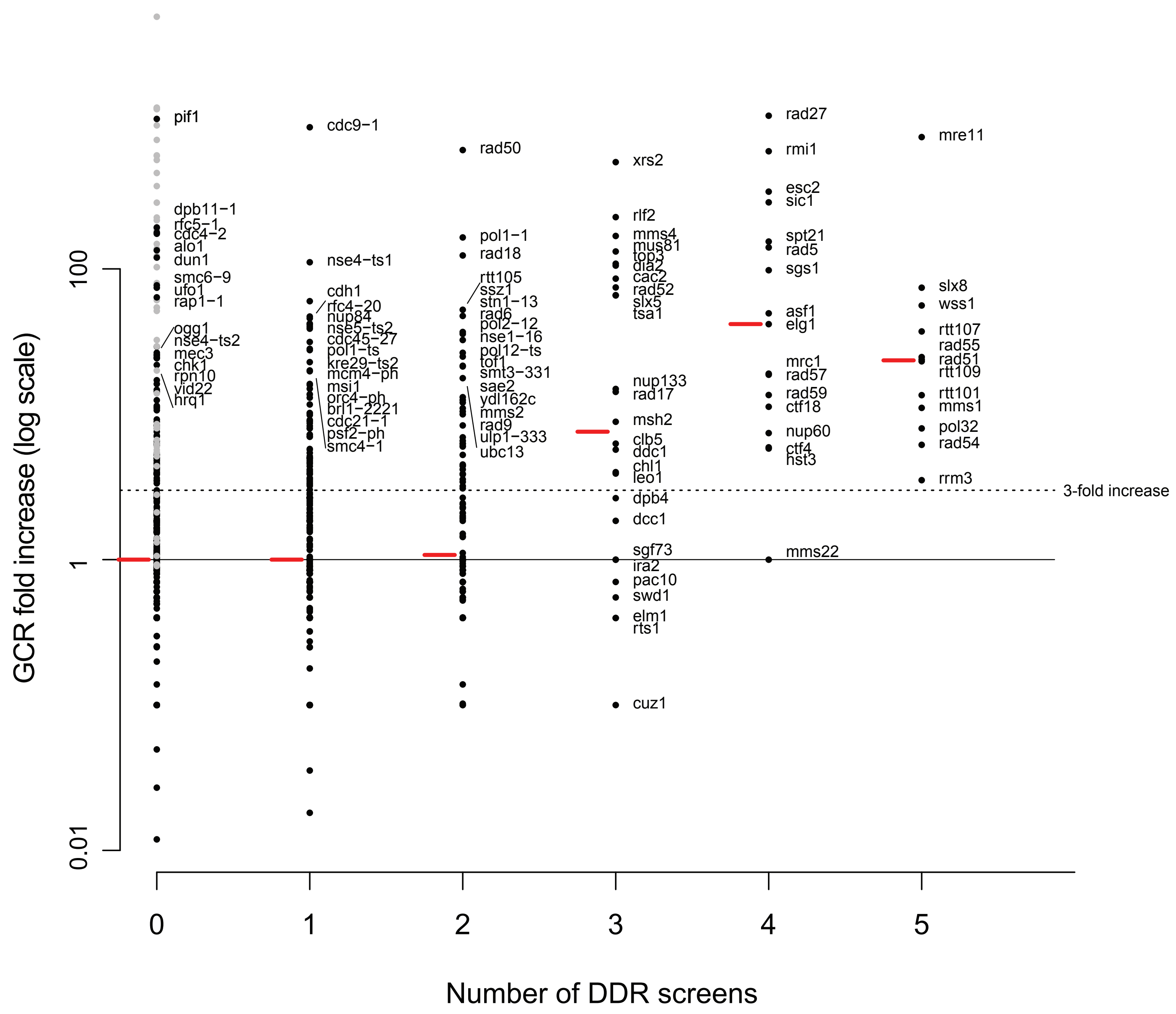
